# Evolutionary stabilisation of stressful metabolism *via* integrated biocomputing and essential-gene metabolic locking circuits

**DOI:** 10.64898/2026.09.01.748751

**Authors:** Zeyu Lu, Craig Barry, Masahiro Tominaga, Anh Phan, Ali Jahanian, Thomas R. Collier, Nicholas A. Wendrick, Samuel Evans, Liam McDonnell, Gabriele Netzel, Tim McCubbin, Shana Jacob, Patrick A. da Roza, Xin Xu, Chengqiang Wang, Jun Ishii, James B. Behrendorff, Alexander S. Beliaev, Claudia E Vickers, Bingyin Peng

## Abstract

Synthetic genetic circuits enable microbial differentiation from growth to production, yet metabolic burden, imbalance and toxicity frequently drive strain degeneration. Here, we use omics analyses and single-cell biosensors to investigate genetic responses in *Saccharomyces cerevisiae* strains engineered with stressful terpene biosynthetic pathways. Under metabolic stress, despite divergent stress responses, cells commonly and progressively lose induction of synthetic genetic circuits. In the prenyl pyrophosphate-accumulating strain, this loss is coupled with activation of amino acid biosynthesis and ribosome biogenesis. Accordingly, the essential glutamine synthetase gene *GLN1* was coupled to induction of heterologous terpene synthesis through integrated two- and three-input biocomputing circuits that enable stringent, patterned ON/OFF switching. The integrated biocomputing and metabolic locking systems not only alleviate production degeneration but also enable non-degenerative adaptation of toxic terpene synthesis, allowing interrogation of evolutionary shifts for further metabolic optimisation.

## Introduction

In multicellular organisms, differentiated cells acquire and maintain specific metabolic patterns that are often genetically stable, with their growth being strictly controlled by natural regulatory mechanisms [1]. In contrast, metabolically engineered microbes frequently undergo strain degeneration when subjected to metabolic burden, toxicity or lethality, resulting in scale-up failure or poor performance during industrial-scale production [2]. Although synthetic approaches are being developed to facilitate differentiation- and development-like transitions in microbial cells by orchestrating genetic regulation and metabolism [3], they have not yet achieved the natural fidelity that enables cells to shift reliably from rapid growth to superior and stable production [4].

In metabolic engineering, synthetic genetic circuit systems have been developed to enable precise control over metabolism and cellular growth [5]. These circuits vary span a range of configurations and functions, including optogenetic pathway activation [6], quorum sensing [6a, 7], and patterned development-like control of cell population [3a, 8]. While genetic circuit designs are advancing towards higher-state decision-making and complex multi-signal regulation [3b, 9], synthetic circuits frequently prove unstable owing to unwanted mutations, genetic escape and evolutionary drift under uncontrolled conditions [2d, 10]. Previously, engineering solutions have primarily focused on improving the overall fitness of synthetic metabolism and genetic circuits [10]. These include the optimisation of the expression profiles of enzyme and regulatory modules, stress-responsive feedback control of productive pathways [11], and strategies that increase regulatory specificity to reduce off-target effects and evolutionary instability [10a]. Fusion of a gene of interest (GOI) and an essential endogenous gene via a ‘leaky’ stop codon-mediated has also been explored, but this approach only stabilised the expression of GOI [2d]. Despite these advances, maintaining hyper-productive and stressful metabolic states remains challenging [4a, 4d, 10c]. The mechanisms by which cells respond and adapt to metabolic toxicity and lethal conditions are still largely unknown [4c, 12]. This knowledge gap largely stems from the lack of sufficiently stringent and evolutionarily stable regulatory tools [5c, 13].

The common budding yeast *Saccharomyces cerevisiae* has been widely used as a chassis organism for biomanufacturing [14] and as a model organism to develop synthetic biology systems for biocomputation [15], data storage [16], as well as understanding genomic evolution and ‘dark’ matters [17]. Here, we explore di- and tri-input circuits to control heterologous terpene synthetic pathways and characterise cellular genetic responses and evolutionary shift upon metabolic imbalance, burden, and toxicity using omics and single-cell biosensors. Through three rounds of design-build-test-learn (DBTL) cycles, we developed an integrated genetic circuit combined with a *GLN1*-mediated metabolic locking system. This system stabilises genetic circuits and facilitates the adaptation of stressful biosynthetic pathways, thereby enabling the interrogation of evolutionary responses to metabolic stresses.

## Results

### Imbalanced prenyl pyrophosphate synthesis inhibits derepression of carbon catabolite repression and activates nitrogen metabolic responses

Accumulation of prenyl pyrophosphates, particularly farnesyl pyrophosphate (FPP), has been hypothesised to cause metabolic stress and cytotoxicity in *S. cerevisiae* strains with an enhanced mevalonate pathway [18]. To investigate this systematically, we constructed a dual-input (auxin + tetracycline) regulatory circuit that independently controls the upstream mevalonate pathway and a downstream FPP-consuming sink (nerolidol synthase AcNES1; Figure 1A). Specifically, *GAL* promoters were used to drive both the mevalonate pathway and AcNES1. Tetracycline induces Gal80 expression, fully repressing the *GAL* promoters and thereby enabling stable strain development and maintenance [5d]. In the absence of tetracycline, auxin triggers degradation of the glucose repressor Mig1, allowing high-level induction of the *GAL* promoters even in the presence of glucose [19]. The strong mevalonate pathway consists of the stabilized Hmg2^K6R^ mutant together with mvaS and mvaE, while the moderate mevalonate pathway contains only Hmg2^K6R^ module. AcNES1 expression drains FPP pools, and strains without AcNES1 accumulate FPP. This 2 × 2 design generated four strains (**Figure 1A-B**) with theoretically graded FPP accumulation under *GAL*-ON conditions: H (high FPP; strong pathway, no AcNES1) > RN (moderate pathway, no AcNES1) > RP (strong pathway + AcNES1) > L (low FPP; moderate pathway + AcNES1).

**Figure 1.**
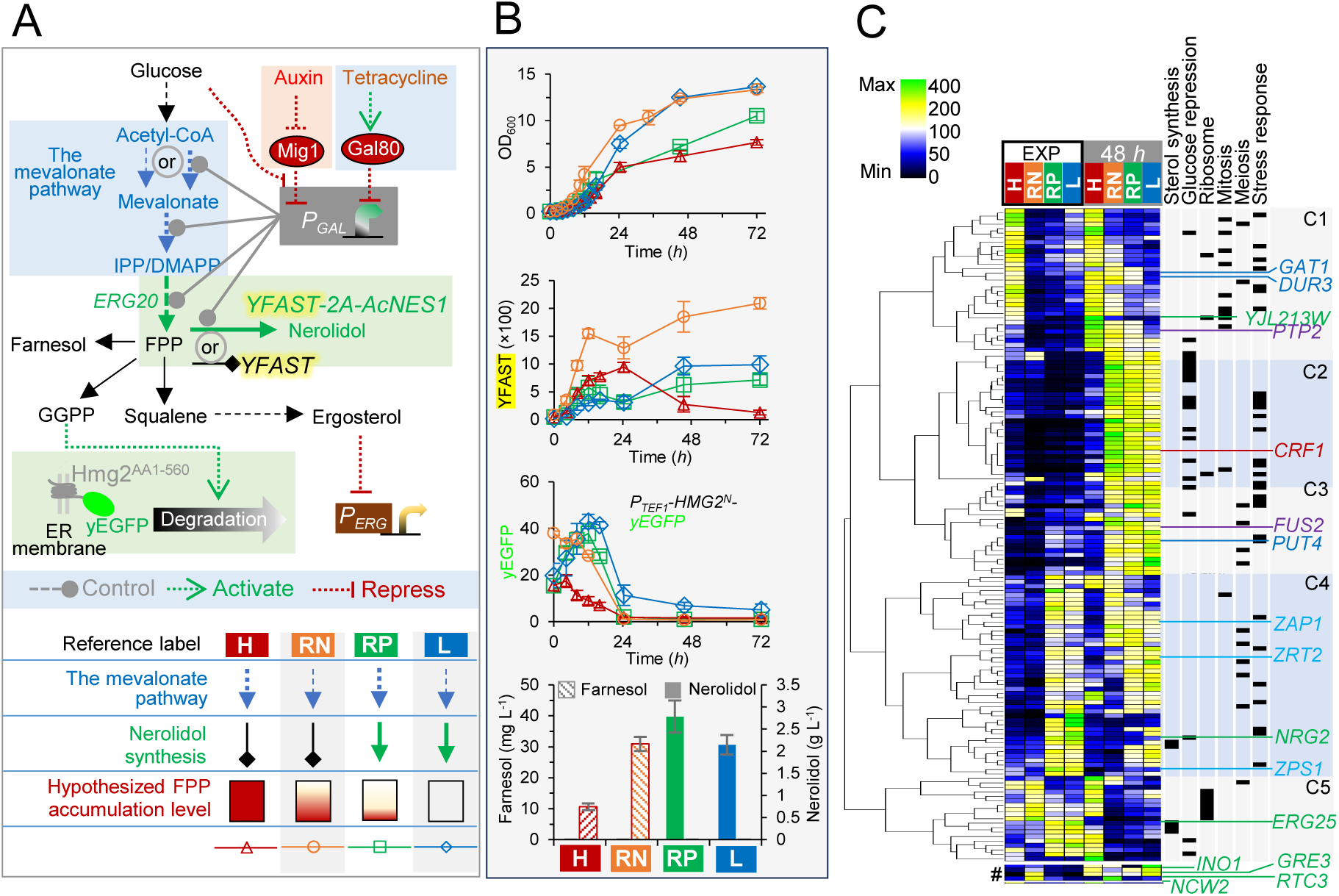
Genetic responses to imbalanced prenyl pyrophosphate synthesis in metabolically engineered yeasts. (**A**) The schematic strain design: metabolic pathways, regulatory circuit logics, and the geranylgeranyl pyrophosphate (GGPP)- and ergosterol-sensing mechanisms. Four engineered strains are: H (ILHA oG57BRB), RN (ILHA oG57H2BRB), RP (ILHA NG57BRB), and L (ILHA NG57H2BRB). (**B**) Characterisation of the four strains in two-phase batch cultivation: growth, YFAST fluorescence, and yEGFP fluorescence, and the titres of nerolidol and farnesol at 72 *h*. (C) Clustering analysis of selected genes in RNA-seq transcriptomics: the genes with the absolute fold-change ≥ 1 (two-tailed *welch*’s test *p*-value ≤ 0.1) in at least three out of four comparisons of H vs. RP and NP vs. L during the exponential growth phase (EXP; OD_600_ = 2.0–2.5) and at 48 *h*. MES-buffered YNB media with 20 g L^-1^ glucose was used in batch cultivation in the presence of 1 mM 1-naphthaleneacetic acid and the absence of tetracycline. ER, endoplasmic reticulum; FPP, farnesyl pyrophosphate; *P_GAL_*, the promoter of the galactose catabolic gene; and *P_ERG_*, the promoter of the ergosterol synthetic gene. Mean values (± standard deviation) are shown (n = 3 biological replicates).

In these four strains, the fluorescence of small fluorescence-activating and absorption-shifting tag (YFAST) [20] served as a direct reporter of *GAL* promoter activity (**Figure 1A**). In parallel, a geranylgeranyl pyrophosphate (GGPP) biosensor was constructed by fusing the N-terminal GGPP-inducible degradation domain of Hmg2 to yEGFP (Hmg2^N^-yEGFP; **Figure 1A**) [21]. Consequently, yEGFP fluorescence inversely reflects intracellular GGPP levels.

These four strains were characterised in batch cultivation under *GAL*-ON conditions (**Figure 1B**). Non-nerolidol-producing strains (H & RN) produced farnesol, a degradation product of FPP, but not nerolidol (**Figure 1B**), consistent with intracellular FPP accumulation. The H strain produced less farnesol than the RN strain, which may be attributable to its poor growth, high cell-death rate (∼ 75% at 48 and 72 *h*; **Supplementary Figure 1**), and reduced *GAL* induction after 48 *h* (YFAST fluorescence; **Figure 1B**). Among the nerolidol-producing strains, the moderate-pathway strain (L) produced less nerolidol than the strong-pathway strain (RP), although both exceeded 2 g L^-1^ nerolidol. The H strain exhibited the lowest Hmg2^N^-yEGFP fluorescence during the exponential growth phase, whereas the L strain maintained the highest fluorescence after 24 *h*. These observations support higher GGPP accumulation in the H strain and a limited GGPP pool in the L strain, reflecting the relative abundance of the upstream precursor FPP.

RNA-seq was performed using Oxford Nanopore sequencing on cells during the exponential growth phase and at 48 *h* in the post-exponential phase (**Figure 1C; Supplementary Figure 2-4**). Clustering analysis revealed decreased mRNA levels of steroid biosynthesis genes (*ERG3*, *ATF2*, *ERG25*, *ERG4*, and *ERG6*) during the exponential growth phase in both the non-nerolidol-producing H and RN strains (**Figure 1C**; Supplementary Figure 2). At 48 *h*, the H strain showed reduced induction of glucose repression target genes (*GAL1/2/7/10* and *MAL32/12/11*; **Figure 1C**; Supplementary Figure 2). In the same strain, several ribosomal genes (*RPL8B/24/2A/5*, *RPA43*, *RPS2*, *RPP0*) were upregulated at 48 h relative to the other strains (**Figure 1C; Supplementary Figure 3**). Mitosis-related genes were also upregulated, whereas meiosis-related genes were downregulated (**Figure 1C**). Genes associated with the responses to zinc starvation (*ZPS1*, *ZRT2*, and *ZAP1*) and various stresses, including DNA replication stress, heat, heavy metals, and drugs, were enriched but exhibited diverse expression patterns (**Figure 1C**).

Transcription factor enrichment analysis indicates that the activators Lys14 and Ifh1 were associated with upregulated genes in the H strain at 48 h (**Supplementary Figure 4**). Although *IFH1* mRNA level was not elevated under these conditions, its paralog *CRF1* was regulated (**Figure 1C**). Ifh1, which regulates ribosomal transcription, and Crf1 function as downstream regulators in the TOR (Target of Rapamycin) signalling pathway [22], which also controls nitrogen and amino acid metabolic genes [23]. Consistent with this, the nitrogen starvation-inducible genes *GAT1* and *DUR3* (but not *PUT4*; **Figure 1C**) and genes involved in amino-acid biosynthesis, including glutamine synthetase (*GLN1*), were upregulated (**Supplementary Figure 5**). In contrast, Hot1, Rtg2, and Yrm1 (a regulator of multidrug resistance) were enriched in association with the downregulated genes in the H strain at 48 *h* (**Supplementary Figure 4**). Clustering analysis showed that *PTP2*, which encodes a phosphotyrosine-specific phosphatase that inactivates the Hot1-upstream activator Hog1 and the *FUS2*-upstream activator Msg5 in the MAPK (Mitogen-Activated Protein Kinase) signalling pathways [24], was upregulated under the same conditions (**Figure 1C**). The reduced activity of Hot1 and Yrm1 may be linked to the diminished induction of the heterologous mevalonate pathway genes controlled by the *GAL* promoters.

Using fluorescent proteins as reporters, we characterised the activities of the *RTC3*, *YJL213W*, *ERG25*, *GRE3*, *NCW2*, and *INO1* promoters (**Supplementary Figure 6**). Only the *ERG25* promoter activities were consistent with the RNA-seq data and showed an inverse association with the hypothesised FPP accumulation. Induction of the *ERG25* promoter was linked to expression of nerolidol synthase (**Supplementary Figure 6G**), suggesting ergosterol starvation in the nerolidol-producing strains. No clear correlation was observed between the cAMP biosensor or cytosolic pH sensor [25] and hypothetical FPP accumulation (**Supplementary Figure 6K-M**). Notably, intracellular pH decreased dramatically in the H strain after entry into the post-exponential phase. In contrast, the RN strain, which carries a moderate mevalonate pathway, appeared to benefit from moderate FPP overproduction, as evidenced by improved growth, higher responses from the *RTC3*, *YJW213W*, *GRE3* promoters, elevated cAMP levels, and higher cytosolic pH in the post-exponential phase (**Supplementary Figure 6**).

### Convergent and divergent physiological responses to engineered sesquiterpene biosynthesis

The balance of prenyl pyrophosphate metabolism can be perturbed at both upstream nodes of the mevalonate pathway and downstream nodes involved in ergosterol and heterologous terpene synthesis. We re-engineered eight versions of the mevalonate pathway (**Supplementary Figure 7**) and evaluated their capacity through nerolidol synthesis. The weakest version (#72) increased nerolidol titre from a background level of 0.1 g L^-1^ to 0.42 g L^-1^ and the strongest version (#6121) enabled production of 2.2 g L^-1^ (**Supplementary Figure 7**). To restrict squalene synthesis, Erg9 was tagged at its C-terminus with an LSSmScarlet-Cln2^PEST^ degradation tag [26]. The Cln2^PEST^ tag reduced LSSmScarlet fluorescence 2.8- to 4-fold compared with the Erg9-LSSmScarlet control (**Supplementary Figure 8**). Tagging Erg9 with LSSmScarlet-Cln2^PEST^ improved nerolidol production by 0.37 ± 0.10 g L^-1^ across five background strains tested (**Supplementary Figure 7**). Disruption of *MIG1* to enable *GAL* promoter induction on glucose further improved nerolidol production by 0.51 ± 0.14 g L^-1^ across all six background strains examined (**Supplementary Figure 7**). In addition, we found that the MvaS A110G mutant [27] proved inferior to the wild-type MvaS in improving nerolidol production. Overexpression of the acetyl-CoA synthase gene *ACS2* in six *mig1Δ* strains yielded no improvement or even a decrease in nerolidol titre (**Supplementary Figure 7**).

To further characterise the stresses associated with terpene metabolic imbalances, we selected six background strains engineered with four different versions of the mevalonate pathway, with or without C-terminal LSSmScarlet-Cln2^PEST^ tagging of Erg9 (**Figure 2A**). These comprised the isogenic background strain carrying version #401 of the mevalonate pathway (previously used in our studies) [18a, 28], the weakest (#72), moderate (#612), and strongest (#6121D; with Erg9-LSSmScarlet-Cln2PEST) versions, as well as two additional strains carrying the Erg9-LSSmScarlet-Cln2^PEST^ tag (#72D and #612D). Into these backgrounds, we introduced the tetracycline-repressible *GAL* circuit, deleted *MIG1*, and expressed additional copy of *GAL* activator *GAL4* gene under the control of the *NRG2* promoter (**Figure 2B**). In the #401 background strain, we examined the induction dynamics of the *GAL* promoter under these engineered genetic regulatory mechanisms. YFAST fluorescence driven by the *SkGAL2* promoter was almost completely repressed when tetracycline was added to both the seed and main cultures, or to the seed culture alone (**Figure 2C**). When tetracycline was omitted from the seed culture, YFAST fluorescence remained severely repressed during the first six hours of the main culture before undergoing rapid induction (**Figure 2C**). Overall, *GAL* promoter-driven YFAST stayed repressed for at least 7.5 cell divisions after tetracycline removal from the medium.

**Figure 2.**
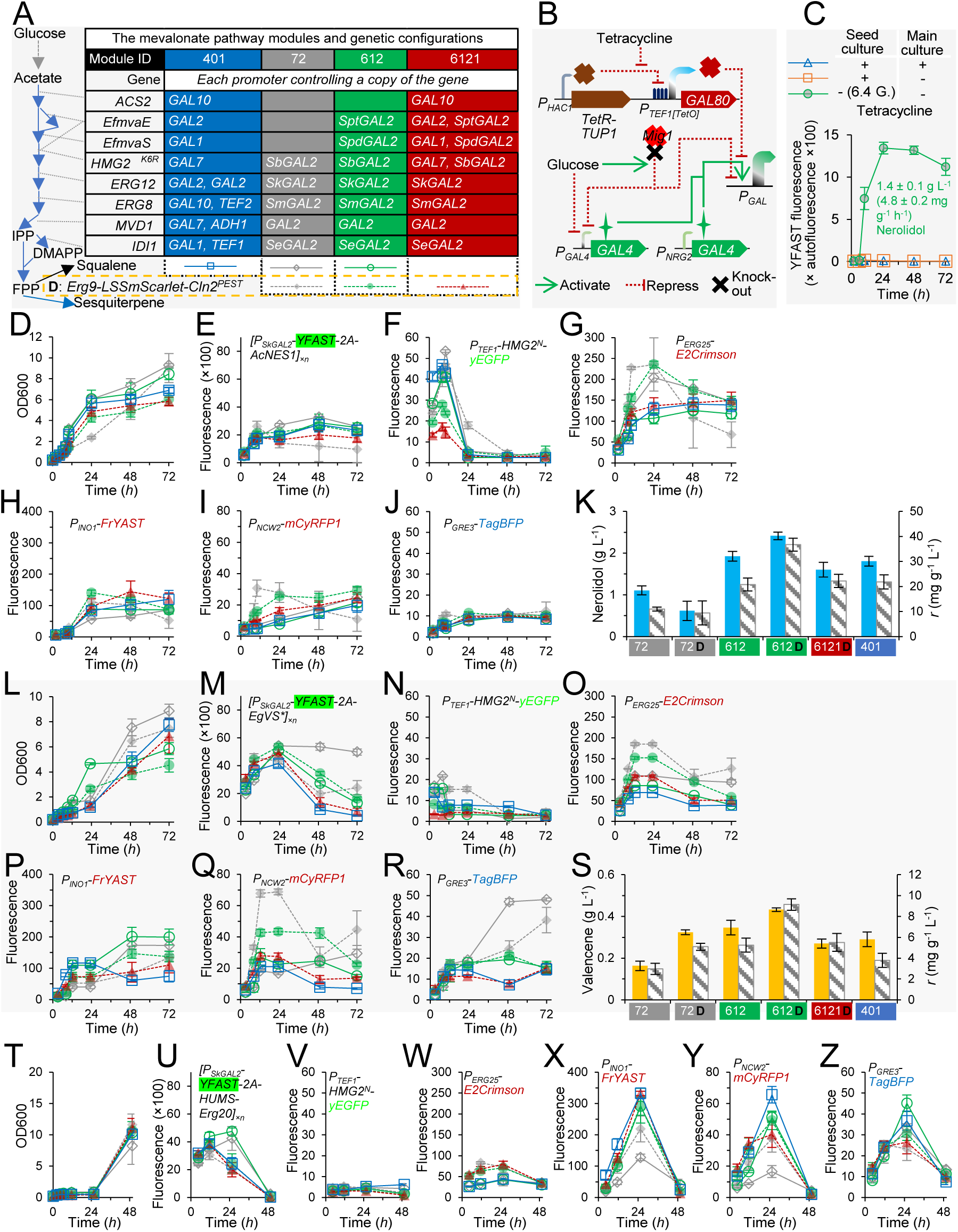
Characterisation of sesquiterpene-producing strains equipped with six versions of farnesyl pyrophosphate (FPP)-supplying pathways and fluorescent reporter responses in batch cultivation. (A) The schematic design of metabolic pathway engineering and the data legend of D-Z. (B) The schematic design of regulatory circuit engineering. (C) Impact of tetracycline addition on YFAST fluorescence induction in a nerolidol-producing strain equipped with the #401 version of the engineered mevalonate pathway. Tetracycline was added or omitted in preculture and main culture. 6.4 G, the average generation time of cell division. (D-K) Characterisation of nerolidol-producing strain. (L-S) Characterisation of nerolidol-producing strain. (T-Z) Characterisation of humulene-producing strain. MES-buffered YNB media with 20 g L^-1^ glucose was used in batch cultivation. Mean values ± standard deviation are shown (n = 3 biological replicates).

Into these background strains we introduced biosynthetic pathways to produce nerolidol, valencene [28c], and humulene [28a] using the multi-copy genome-integrating HapAmp vector [28b]. In addition, six fluorescent reporters were incorporated to monitor intracellular genetic and metabolic responses: YFAST for *GAL* promoter activity, Hmg2^N^-yEGFP for GGPP levels, E2-Crimson for the *ERG25* promoter, FrFAST for the *INO1* promoter, mCyRFP1 for the *NCW2* promoter, and TagBFP for the *GRE3* promoter. This yielded three groups of strains: nerolidol-producing (**Figure 2G–2K**), valencene-producing (**Figure 2L–2S**), and humulene-producing (**Figure 2T–2Z**). Each group comprises six strains. All 18 strains were tested under *GAL*-ON conditions in batch cultivation and exhibited both convergent and divergent genetic and metabolic responses.

In both nerolidol- and valencene-producing strains, the moderate mevalonate pathway combined with Erg9-LSSmScarlet-Cln2^PEST^ (#612D) yielded the highest titres, whereas the strongest mevalonate pathway version (#6121D) resulted in lower production (**Figure 2K** & **2S**). Consistent with pathway design, Hmg2^N^-yEGFP fluorescence was lower in the #6121D strains than in the #612D strains during the exponential growth phase (**Figure 2F & 2N**). Overall, Hmg2^N^-yEGFP fluorescence was 4.3 ± 2.9-fold lower overall in the valencene-producing strains than in the isogenic nerolidol-producing strains (**Figure 2F** & **2N**). Higher YFAST fluorescence in the valencene-producing strains (**Figure 2E** & **2M**) may indicate higher expression of terpene synthase, as YFAST and the sesquiterpene synthase are translated from the same transcript. Together, these results suggest that nerolidol synthase (AcNES1) has superior catalytic properties relative to valencene synthase (EgVS*) [13] and that the sesquiterpene synthase step constitutes a major bottleneck for efficient heterologous sesquiterpene production.

Erg9-LSSmScarlet-Cln2^PEST^ tagging improved nerolidol production in the #612 background strain but not in the weakest #72 background strain (**Figure 2K**), while enhancing valencene production in both backgrounds (**Figure 2S**). The Erg9-LSSmScarlet-Cln2^PEST^ strains (#72D & #612D) exhibited higher E2-Crimson fluorescence driven by the *ERG25* promoter than isogenic *ERG9*-wildtype strains (#72 & #612; **Figure 2G & 2O**). This indicates that flux restriction at the Erg9 node triggers ergosterol starvation. Consistent with this, Erg9-LSSmScarlet-Cln2^PEST^ modification reduced the specific maximal growth rates in both nerolidol- and valencene-producing strains (**Supplementary File 2**). Decreased GGPP-sensing Hmg2^N^-yEGFP fluorescence further supports that constraining Erg9 nodes increases prenyl pyrophosphate pools. In addition to these positive effects, restricting Erg9 negatively affected on nerolidol production in the #72 background strain, which may result from more severe ergosterol starvation in this strain.

All six humulene-producing strains exhibited a lag phase for at least 24 hours in the main culture, followed by rapid growth between 24 and 48 h (**Figure 2T**). However, the recovered cells had lost YFAST fluorescence, indicating that unknown mechanisms caused repression of the *GAL* promoter (**Figure 2U**). In these strains, Hmg2^N^-yEGFP fluorescence remained at basal levels throughout cultivation (**Figure 2V**), indicating substantial GGPP accumulation. Although E2-Crimson fluorescence was higher in the Erg9-LSSmScarlet-Cln2^PEST^ strains than in the *ERG9* wild-type strains, it was not induced to the levels observed the nerolidol- and valencene-producing strains (**Figure 2W**). Collectively, these findings indicate that the humulene-producing strains experienced severe metabolic imbalances, leading to substantial cellular stress and subsequent strain degeneration.

FrFAST, mCyRFP1, and TagBFP fluorescence, driven by the *INO1*, *NCW2*, and *GRE3* promoters, respectively, were simultaneously induced during the lag phase in humulene-producing strains, independent of the designed capacity of the mevalonate pathway (Figure 2X-Z). However, in nerolidol- and valencene-producing strains, their responses did not show a consistent pattern (**Figure 2H–J and 2P–R**). These results demonstrate that stress responses in strains producing different sesquiterpenes are highly diverse and influenced by multiple complex factors.

We also constructed and characterised six isogenic strains lacking a heterologous terpene synthase. These strains displayed growth and fluorescent reporter profiles (**Supplementary Figure 10**) that differed distinctly from those of the nerolidol-, valencene-, and humulene-producing strains. These results further support that producing different sesquiterpenes contributes to diverse genetic responses observed. Notably, the #72 and #72D strains without heterologous sesquiterpene synthesis exhibited exponential growth with a μmax of approximately 0.21 h^-1^, indicating that the severe stress effects seen in the isogenic humulene-producing strains were specifically linked to humulene production.

We further analysed the population heterogeneity of the best nerolidol- and valencene-producing strains (#612D) by examining the distribution of YFAST fluorescence (**Supplementary Figures 11-13**). The proportion of cells that lost YFAST fluorescence increased continuously over time (**Supplementary Figures 11 and 12**), reaching 44 ± 14% in the nerolidol-producing strain and 63 ± 4% (**Supplementary Figure 13A**) in the valencene-producing strain at 72 *h*. Importantly, the nerolidol-producing cells remained propidium iodide-negative (**Supplementary Figure 13B**), indicating that the loss of YFAST fluorescence (*GAL* induction) was not caused by cell death but by other intracellular genetic responses.

### De-repression of carbon catabolite repression is essential for monoterpene limonene production

We then examined the monoterpene (limonene)-producing strains, which were negatively impacted by FPP overproduction due to the presence of native *ERG20* [21, 29]. We first selected the #401 background strain to re-evaluate the effects of *ERG20* downregulation on limonene production. We used the *ERG25* promoter to drive expression of the endogenous *ERG20* gene, modified with an N-terminal N-end-rule degron tag (K3K15, KN113, KN119; **Figure 3A**) [18a], in the #401 background strain with *mig1Δ* and wildtype-*ERG4* only. The limonene biosynthetic module was introduced using the multi-copy genome-integrating HapAmp vector [28b]. An isogenic *ERG20*-wildtype strain was also constructed for the reference. Surprisingly, all four strains produced limonene at unprecedented titres from 20 g L^-1^ glucose in batch culture, with the *ERG20*-wildtype strain even reaching 1.9 g L^-1^, the K3K15 strain reaching 2 g L^-1^, and the KN113 and KN119 strains producing slightly lower titres (**Figure 3B**). YFAST fluorescence was induced during the exponential growth phase in these strains, and the KN113 strain was superior in growth (**Supplementary Figure 14A & 14B**). Notably, the monoterpene titre was doubled relative to our previous best strain that also utilised the #401 version of the mevalonate pathway [28b]. Further verification confirms that *mig1Δ* is critical for limonene production in the *ERG20*-wildtype background (**Figure 3B**) and induction of YFAST fluorescence driven by the *GAL* promoter (**Supplementary Figure 14D**).

**Figure 3.**
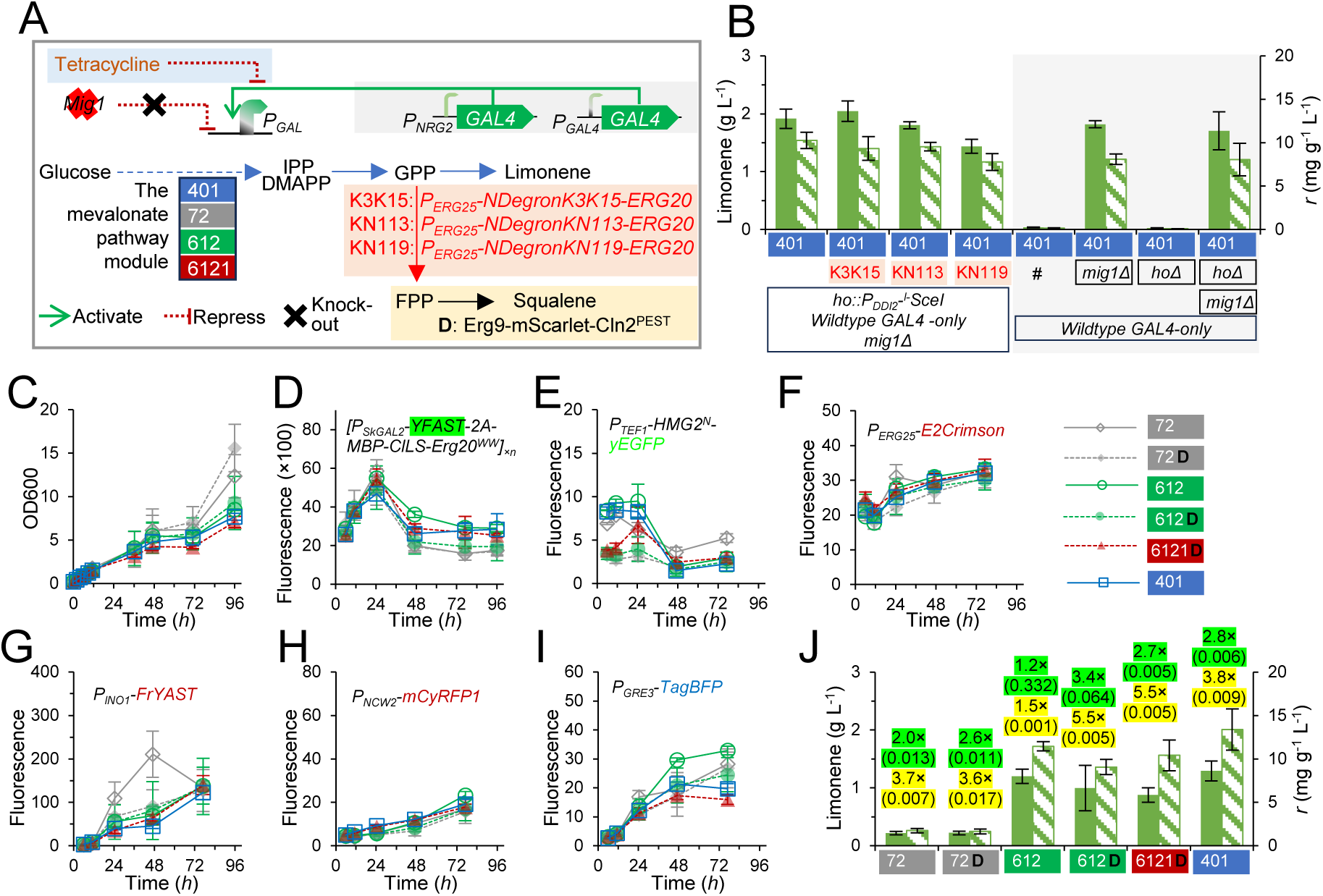
Characterisation of monoterpene limonene-producing strains. (A) The schematic design of the metabolic pathway and regulatory circuit engineering. (B) Impact of *MIG1* disruption on limonene production. Growth profiles are shown in Supplementary Figure 13. (C-J) Characterisation of limonene-producing strains with farnesyl pyrophosphate (FPP) synthase fused with an N-terminal K3K15 degron. The values above the bars in J are the fold-change values and two-tailed *Welch*’s test *p*-values (in brackets) of limonene titre (green shadow) and specific production rate (yellow shadow) compared to the isogenic *ERG20*-wildtype strains. MES-buffered YNB media with 20 g L^-1^ glucose was used in batch cultivation. Mean values ± standard are deviation shown (n = 3 biological replicates).

We characterised twelve (6×2) monoterpene-producing strains, which were engineered similarly to above nerolidol/valencene/humulene-producing strains (*mig1Δ* + *P_NRG2_-GAL4*) with wildtype genomic *ERG20* (**Supplementary Figure 15**) or genomic *P_ERG25_-NdegronK3K15-ERG20* modification (**Figure 3C-J**). The *P_ERG25_-NdegronK3K15-ERG20* modification improved both limonene titre and specific production rate in all six background strains. While the improvement was moderate in the #612 background, it resulted in at least a 2-fold increase in the other five backgrounds (**Figure 3J**). In the *ERG20* wild-type background, the #401 strain only produced 0.46 g L^-1^ (**Supplementary Figure 15**), suggesting that constitutive high-level induction of the synthetic pathway is not tolerated. In contrast, the #612 strain outperformed the others in both *ERG20*-wildtype and *P_ERG25_-NdegronK3K15-ERG20* strains. Erg9-LSSmScarlet-Cln2^PEST^ yielded no or negative effects on limonene production (**Figure 3J & Supplementary Figure 15H**).

Notably, the *P_ERG25_-NdegronK3K15-ERG20* modification did not eliminate farnesol production; instead, all six strains produced farnesol at comparable levels (**Supplementary Figure 16**). This modification increased farnesol production in the #401 strain, likely by alleviating metabolic stress, as evidenced by improved induction of *GAL* promoter-driven YFAST fluorescence (**Figure 3D** & **Supplementary Figure 15B**).

The limonene-producing strains did not exhibit dramatic responses in E2-Crimson, FrFAST, mCyRFP1, or TagBFP fluorescence (**Figure 3G-I**). However, similar to the sesquiterpene-producing strains, Cells gradually lost YFAST fluorescence after 24 h, eventually reaching 19 ± 9% at 96 *h* (**Supplementary Figures 17 & 18**). Strikingly, cell agglomeration appeared in these limonene-producing strains during batch cultivation (see example in **Supplementary File 3**).

### Modular design of an integrated biocomputing circuit and metabolic locking system

Metabolic stress arising from constitutively induced heterologous terpene biosynthesis caused pronounced population heterogeneity, most notably the loss of *GAL* induction in a substantial subpopulation or the entire population in humulene-producing strains. Furthermore, transferring nerolidol-producing cells from a 72-*h* ‘*GAL*-ON’ culture into fresh medium failed to restore *GAL* induction; instead, the entire population lost YFAST fluorescence (**Supplementary Figure 19**). The loss of *GAL* induction may reflect the inherent flexibility and plasticity of both natural and synthetic genetic regulatory networks. Based on the RNA-seq and single-cell reporter data, we designed di- and tri-input regulatory (biocomputing) circuits (**Figure 4A**: ①, ②, & ③) together with an integrated *GLN1*-mediated metabolic locking system (**Figure 4A**: ④) to address these issues.

**Figure 4.**
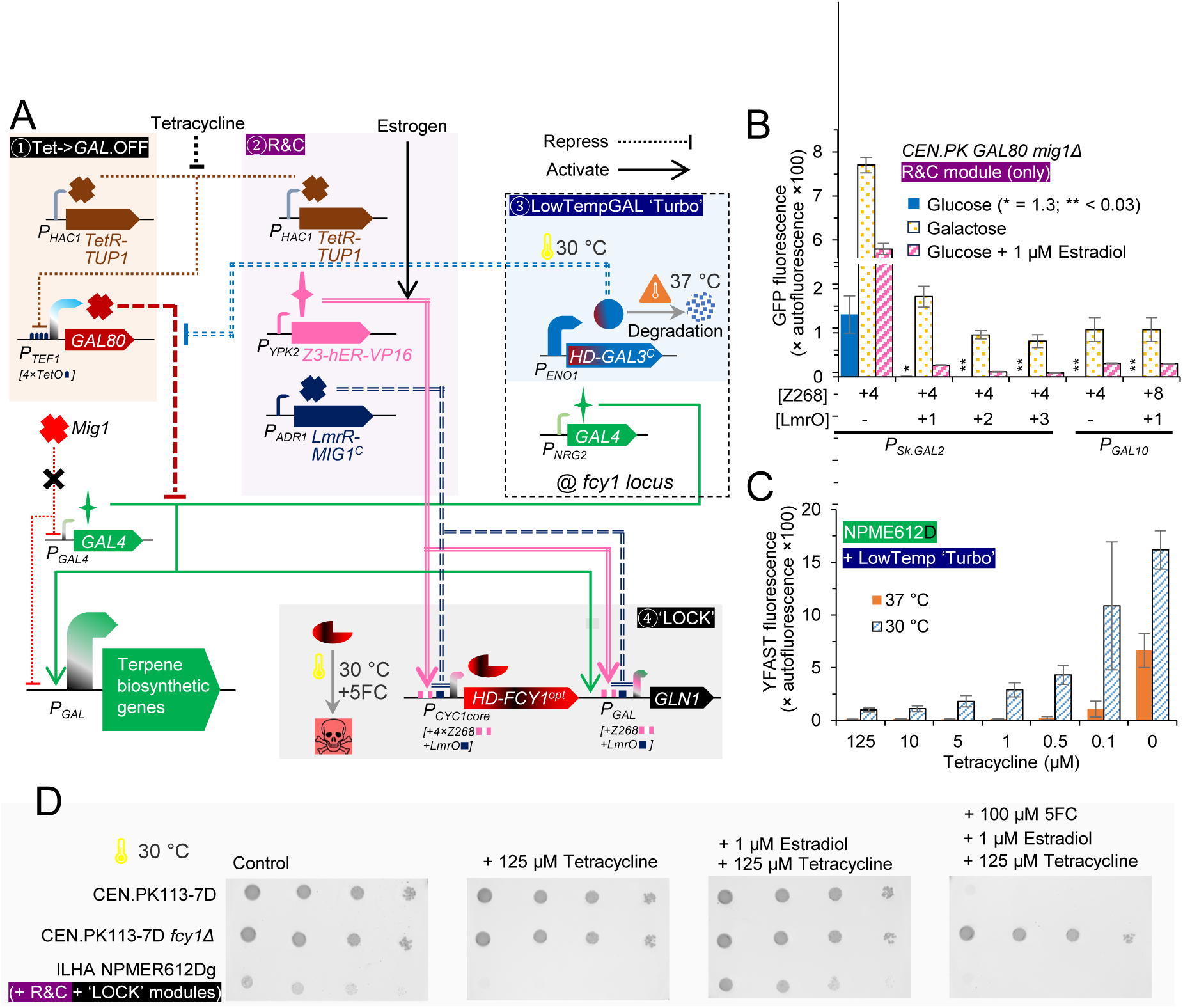
Modules of regulatory (biocomputing) circuits and the *GAL* promoter-stabilising metabolic locking mechanism. (A) The design of regulatory circuits: the tetracycline-mediated *GAL* repression circuit module, the redundant and complementation module (R&C), the low-temperature-inducible (LowTemp) *GAL* ‘Turbo’ module, and the *GLN1*-mediated metabolic locking module. (B) Characterisation of the galactose|estrogen ‘OR’ logic gate for induction of the hybrid *GAL* promoters. (C) Characterisation of the LowTempGAL ‘turbo’ module on tetracycline-mediated *GAL* repression at 30 and 37 °C. (D) Validation of growth phenotypes of the cells equipped with Tet->*GAL*.OFF, R&C, and ‘LOCK’ modules. Mean values ± standard deviation are shown (n = 3 biological replicates; B-C). n = 1 in D.

We postulated that the potential leakage from the hybrid *TEF1-TetO* promoter could permit low-level expression of *GAL80* (**Figure 4A**: ①), thereby contributing to *GAL* repression. To counteract this, we introduced a second copy of the *TetR*-*TUP1* expression cassette as a redundant repressor (**Figure 4A**, ②).

RNA-seq analysis of the H strain showed that diminished de-repression of glucose repression genes was accompanied by upregulation of amino-acid biosynthesis genes, activation of the nitrogen starvation response (*GAT1* and *DUR3*), and increased ribosomal biogenesis via TOR-downstream activator Ifh1 [22]. We therefore hypothesised that placing an essential amino acid biosynthetic gene under the *GAL* promoter would render *GAL* induction obligatory for cell viability. From the amino-acid biosynthesis genes (**Supplementary Figure 5**), we selected glutamine synthetase (*GLN1*) for the following reasons: (1) *GLN1* is present as a single copy in *S. cerevisiae* genome; (2) glutamine is the primary nitrogen donor for the biosynthesis of many other amino acids; and (3) glutamine functions as a key signalling molecule in nitrogen catabolite repression and TOR signalling [30]. Consequently, *GLN1* was placed under *GAL* promoter control to create a metabolic locking module (**Figure 4A**, ④)

Because a strain carrying *GAL*-driven *GLN1* as its sole copy cannot be maintained under *GAL*-OFF conditions, we designed an estrogen-inducible complementation circuit (**Figure 4A**: ②). This circuit employs the artificial trans-activator Z3-hER-VP16 (a fusion of a Z268-binding zinc finger domain, the human Estrogen Receptor, and the VP16 activation domain) to drive a hybrid *GAL-*[Z268] promoter in the presence of estrogenen (**Figure 4A**: ④). To suppress leakiness of the estrogen-inducible system, we previously incorporated the artificial repressor LmrR-Mig1^C^ (**Figure 4A**: ②) [31]. Initial tests of a hybrid *SkGAL2*-[4×Z268] promoter showed residual leakage in the *mig1Δ* background; inserting an LmrO motif eliminated this leakage but reduced induction strength (**Figure 4B**). Two hybrid *GAL10*-[Z268] promoters, however, exhibited no detectable basal expression even without LmrR-Mig1^C^ repression (**Figure 4B**).

To further guard against potential malfunction of Z3-hER-VP16 or LmrR-Mig1^C^, we added a 5-fluorocytosine counter-selection module. In this design, a hybrid *CYC1*core-[4×Z268+LmrO] promoter drives a codon-optimised fusion of a heat-inducible degron and cytosine deaminase (**Figure 4A**: ④).

To validate the stringency of the complete mechanism, we constructed a nerolidol-producing #612D strain (ILHA NPMER612Dg) carrying the full Redundancy-&-Complementing (R&C) and metabolic locking ‘LOCK’) modules (Figure 4A), with the sole *GLN1* copy under the control of the *GAL10*-[4×Z268] promoter. The dot-dilution assays confirmed that tetracycline abolished growth, estrogen restored growth, and 5-fluorocytosine was lethal under estrogen induction (**Figure 4D**).

Finally, we tested the compatibility of the low-temperature-inducible *GAL* (‘LowTempGAL’) ‘turbo’ module (**Figure 4A**: ③) [28a]. Expressing a heat-inducible-degron-Gal3^C^ fusion produced limited leakage from the *GAL* promoter in the presence of tetracycline at 30 °C (**Figure 4C**). This module therefore provides both an additional route to ensure *GAL* induction and a second, estrogen-independent mechanism for *GLN1* expression.

### *GLN1*-mediated metabolic locking enables adaptation to sesquiterpene biosynthesis stress

#### Adaptation of nerolidol-producing strain

To validate whether *GLN1*-mediated metabolic locking mechanism can stabilise *GAL* induction, we adaptively cultivated the strain NPMER612Dg, which harbours the Redundancy & Complementation circuit (Figure 4: ② R&C) and the metabolic locking module (Figure 4: ④ ‘Locking’). This strain was developed and maintained in the medium containing tetracycline and estrogen. To initiate the culture in the absence of tetracycline and estrogen, the preculture was supplemented with 1 g L^-1^ glutamine to complement the glutamine deficiency and allow sufficient cell divisions to dilute residual Gal80p, thereby enabling *GAL* induction. In subsequent cultures (without dodecane overlay), glutamine was omitted. Strain NPMER612Dg maintained the binary YFAST fluorescence induction profiles in the second and third transferring cultures (**Figure 5A-B**), indicating that *GLN1*-mediated metabolic locking can sustain *GAL* induction. The final culture in transferring cultivation (**Figure 5A-B**) was subsequently grown in the presence of tetracycline and estrogen to prepare a glycerol stock of adapted strain ILHA NPMER612DgE.

**Figure 5.**
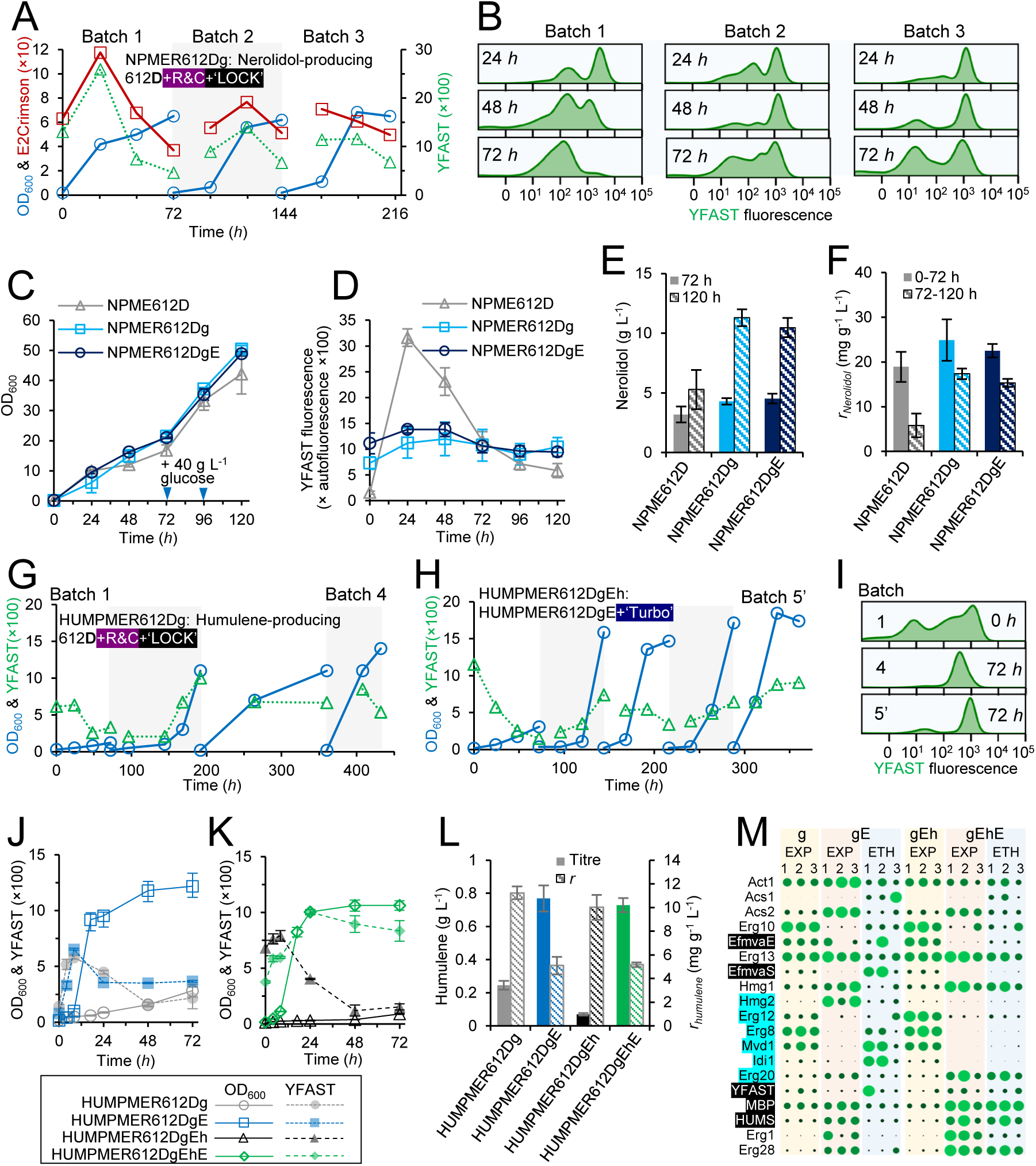
Integrated biocomputing and *GLN1*-mediated metabolic locking mechanisms enabling the adaptation of nerolidol-producing (A-F) and humulene-producing (G-M) yeasts under metabolic stresses. (A) Transferring cultivation of nerolidol-producing strain NPMER612Dg equipped with the redundant and complementation circuit module (R&C) and the metabolic locking (‘LOCK’) module in MES-buffered YNB broth with 20 g L^-1^ glucose as the carbon source. (B) Density of YFAST fluorescence in cells sampled in transferring cultivation in A. (C-F) Semi-fed-batch fermentation of the nerolidol-producing #612D strain (NPME612D), NPMER612Dg, and the adapted strain NPMER612DgE: growth, YFAST fluorescence, nerolidol titre, and the specific nerolidol production rate (*r*). The MES-buffered 4×YNB broth with 40 g L^-1^ glucose as the initial carbon source. Glucose (40 g L^-1^) was fed at 72 *h* and 96 *h*. (G) Transferring cultivation of the humulene-producing strain HUMPMER612Dg equipped with the R&C and ‘LOCK’ modules in MES-buffered YNB broth with 20 g L^-1^ glucose as the carbon source. (H) Transferring cultivation of the derivative of the adapted strain HUMPMER612DgE in which the low-temperature-inducible *GAL* ‘Turbo’ (heat-inducible GAL3^C^) module was introduced. MES-buffered 4×YNB broth with 40 g L^-1^ glucose was used. (I) Density of YFAST fluorescence in cells sampled in transferring cultivation in G & H. (J-L) Batch cultivation of humulene-producing strains in MES-buffered YNB broth with 20 g L^-1^ glucose as the carbon source. (M) Relative abundance of the enzymes in terpene synthetic pathways (data are presented in Supplementary Figure 27 using different scales). The circle radius represents the abundance normalised in the range of 0-100. Enzymes in black background and white font were expressed under the control of the *GAL* promoter solely. Enzymes in cyan background were expressed with the additional heterogenous copy driven by the *GAL* promoter. Mean values ± standard deviation are shown (n = 3 biological replicates; C-F, J-L).

We next characterised the strains NPME612D (nerolidol-producing #612D from **Figure 2**), NPMER612Dg, and NPMER612DgE in semi-fed-batch cultivation with a dodecane overlay (**Figure 5C-F**). The cultures were initiated with 40 g L^-1^ glucose, and two additional pulses of 40 g L^-1^ glucose were fed at 72 *h* and 96 *h*. Strain NPME612D exhibited hyperactive *GAL* induction at 24 *h* and 48 *h* (**Figure 5D & Supplementary Figure 20A**) and produced 3.2 g L^-1^ nerolidol at 72 *h* (**Figure 5E**). At the single-cell level, most of the population had lost the YFAST fluorescence at 72 *h* (51 ± 7%), with further loss thereafter (**Supplementary Figure 20A**). This loss of induction coincided with a marked decline in overall productivity (**Figure 5E-F**).

In contrast, both the strain carrying the *GLN1*-mediated locking mechanism (NPMER612Dg) and its adapted derivative (NPMER612DgE) maintained stable YFAST fluorescence and sustained nerolidol production after 72 *h* (**Figure 5D-F**). Notably, YFAST fluorescence did not reach the high levels observed in the strain NPME612D (**Figure 5D & Supplementary Figure 20A**). At the single-cell level, the unadapted strain NPMER612Dg exhibited heterogeneous YFAST fluorescence patterns across the three biological replicates during the first 72 *h*; these patterns gradually became more uniform, with only 14 ± 3 % of cells having lost YFAST fluorescence by 120 *h* (**Supplementary Figure 20B**). The adapted strain NPMER612DgE, however, showed highly consistent and constitutive induction of YFAST fluorescence in the major population throughout the entire cultivation period (**Supplementary Figure 20C**).

#### Adaptation of humulene-producing strain

We next focused on adapting humulene-producing strain using the *GLN1*-mediated metabolic locking mechanism. In the presence of the Redundancy & Complementation circuit (Figure 4: ② R&C) and the metabolic locking module (Figure 4: ④ ‘Locking’), the humulene-producing strain (ILHA HUMPMER612Dg) exhibited a prolonged 150-*h* lag phase during which YFAST fluorescence remained low (**Figure 5G**). After 150 *h*, growth gradually resumed upon serial transfers, accompanied by the emergence of a population displaying homogeneous YFAST fluorescence (**Figure 5I**). Cells from the final culture were grown in the presence of tetracycline and estrogen to generate glycerol stocks of the adapted strain ILHA HUMPMER612DgE.

We initially planned for semi-fed-batch cultivation for HUMPMER612DgE in 4×YNB medium with 40 g L^-1^ glucose. However, YFAST fluorescence in this strain remained in both the preculture (−tetracycline, + glutamine) and the main culture (-tetracycline, - glutamine). To enhance *GAL* induction, we introduced the LowTempGAL ‘Turbo’ construct (**Figure 4A**: ③) into HUMPMER612DgE, to generate the strain ILHA HUMPMER612DgEh. This modification permits limited constitutive *GAL* induction at 30 °C (**Figure 4C**), thereby enabling growth in the absence of estrogen and glutamine. In 4×YNB medium with 40 g L^-1^ glucose, HUMPMER612DgEh exhibited a 120-*h* lag phase, followed by adaptive evolution through serial transfers, yielding the final adapted strain ILHA HUMPMER612DgEhE (**Figure 5H**).

The four humulene-producing strains (HUMPMER612Dg, HUMPMER612DgE, HUMPMER612DgEh, and HUMPMER612DgEhE) were subsequently evaluated in batch cultivation. The unadapted strains (HUMPMER612Dg and HUMPMER612DgEh) showed poor growth (**Figure 5J-K**) and produced only ∼0.25 and ∼0.05 g L^-1^ humulene, respectively, at 72 *h* (**Figure 5L**). In contrast, the adapted strains (HUMPMER612DgE and HUMPMER612DgEhE) displayed typical exponential growth (**Figure 5J-K**) and reached significantly higher titres of ∼0.77 and 0.73 g L^-1^ humulene at 72 *h* (**Figure 5L**), representing more than 60% improvement relative to our previous humulene-producing strain [28a]. However, the adapted strains exhibited approximately half the specific humulene production rate of the unadapted strains (**Figure 5L**), indicating that adaptation came at the expense of specific productivity.

We also attempted fed-batch cultivation of the adapted humulene-producing strains HUMPMER612DgE and HUMPMER612DgEhE (**Supplementary Figure 21**). Glucose feeding alone at 72 *h* did not support further growth. Supplementation with glucose together with yeast extract and peptone broth enabled a second growth phase in HUMPMER612DgE, but the specific humulene productivity was compromised (**Supplementary Figure 21**). HUMPMER612DgEhE showed only a slight increase in OD_600_ with no substantial improvement in humulene titre (**Supplementary Figure 21**).

#### Proteomics analysis of *GLN1*-mediated metabolic locking and adaptive evolution

We also characterised valencene- and limonene-producing strains carrying the *GLN1*-mediated metabolic locking mechanism without prior cultivation in fed-batch mode **(Supplementary Figure 22-23**). Similar to the nerolidol-producing strain, the valencene-producing strain lacking the *GLN1* locking mechanism exhibited hyperactive YFAST induction followed by loss of induction in most of the population (**Supplementary Figure 22**). Introduction of the locking mechanism resulted in significantly slower growth and diminished YFAST induction (**Supplementary Figure 22**). The limonene-producing strain carrying the locking mechanism maintained relatively stable limonene production, with productivity comparable to that observed in batch cultivation (**Supplementary Figure 23 & Figure 3**). However, a substantial proportion of cells lost YFAST fluorescence after 72 *h*, reaching 43 ± 5% at 120 *h* (**Supplementary Figure 23**). Propidium iodide staining revealed a lethality rate of 62 ± 7% at 120 *h* (**Supplementary Figure 24**), indicating that cell death was the primary cause of YFAST fluorescence loss in the subpopulation.

To investigate the genetic and physiological changes associated with terpene biosynthesis, *GLN1*-mediated metabolic locking and adaptive cultivation, we performed proteomic analysis of the nerolidol-, humulene-, valencene- and limonene-producing strains together with the wild-type CEN.PK strain (**Supplementary Figure 25-28**). Strain producing different terpene products exhibited significantly divergent proteomic responses. *GLN1*-mediated metabolic locking increased the abundance of the mevalonate pathway enzymes (mvaS, mvaE, Erg12, Erg8, Mvd1, Idi1, and Erg20) in nerolidol- and valencene-producing strains, which was consistent with its *GAL* stabilisation effects (**Supplementary Figure 27**). Notably, glycerol/dihydroacetone metabolic enzymes (Gcy1, Dak2, and Gpp2) were significantly upregulated in the unadapted *GLN1*-locked valencene- and humulene-producing strains, and were downregulated after adaptation in the humulene-producing strains (**Supplementary Figure 27**).

Notably, adaptation of the humulene-producing strains was accompanied by a dramatic decrease in the abundance of the mevalonate pathway enzymes (**Figure 5M & Supplementary Figure 27**). The biological replicates of HUMPMER612DgE exhibited highly diversified adaptive patterns (**Figure 5M**). In contrast, HUMPMER612DgEhE showed a highly consistent adaptive pattern across three biological replicates, characterised by a dramatic decrease in the abundance of all enzymes in the engineered mevalonate pathway and a concomitant increase in the abundance of humulene biosynthetic enzymes (**Figure 5M**). Furthermore, the abundance of all enzymes in the pentose phosphate pathway decreased in the adapted strains (**Supplementary Figure 27**). These proteomic changes not only explain the reduced humulene productivity but also suggest that further optimisation should decrease flux push through the mevalonate-pathway module and increase flux drain through humulene synthase.

## Discussion

Degeneration of synthetic phenotypes, including reduced productivity and loss of genetic reporter outputs, remains a common challenge in metabolic engineering and synthetic biology [5c, 14a]. This phenomenon is largely driven by the selective disadvantage of slowed growth under metabolic burden. In this study, we challenged yeast cells with synthetic metabolic stresses arising from terpene biosynthesis and observed both divergent genetic responses among strains engineered to produce different terpenes and a convergent progressive loss of *GAL* promoter activity, either in subpopulations or across the entire population. Guided by transcriptomic data and principles of biocomputation and molecular evolution, we developed a *GLN1*-mediated metabolic locking mechanism that stabilises *GAL* induction, thereby enabling cellular adaptation toward more balanced metabolism and sustained production.

We observed strong metabolic stress effects that caused rapid strain degeneration. These effects can be attributed to strong pathway induction, pathway imbalance, and product toxicity. The *mig1Δ* genotype emerged as an important contributing factor in the nerolidol- and humulene-producing strains, as comparable degeneration was not observed in isogenic *MIG1* strains under strong *GAL* induction [28a]. In contrast, *mig1Δ* proved beneficial for limonene production by alleviating the negative effects of metabolic imbalance. The severe phenotype of humulene-producing strains appears linked to product accumulation itself: isogenic strains lacking the humulene synthase displayed normal growth and *GAL* induction (−72 and −72D in **Supplementary Figure 9**), whereas the corresponding humulene producers suffer pronounced burden.

The precise molecular mechanisms of stress sensing, signal transduction, and subsequent repression of the synthetic *GAL* circuit remain largely unclear, except for components of the sterol response pathway (for example, the *ERG25* promoter and Hmg2^N^) [32]. Prenyl pyrophosphate accumulation was associated with cell death and with altered transcriptional programmes in surviving cells, including signatures consistent with TOR-pathway involvement (Ifh1/Lys14 enrichment and induction of *GAT1* and *DUR3*). Nevertheless, both biosensor and omics data show that strains producing different products and pathway configurations mount divergent genetic responses, yet the complete or partial loss of *GAL* induction is the common feature linked to degeneration.

The *GLN1*-coupling mechanism effectively stabilises *GAL* induction and thereby sustains production of heterologous terpenes. Different from previous product-specific biosensor-based metabolic coupling systems [12b, 33], this work demonstrates a generalisable engineering strategy that exploits gene essentiality to stabilise genetic circuits. In the final design we incorporated repressor redundancy (additional TetR-Tup1 copy) and Gal3^C^-mediated functional repression of Gal80. We could not distinguish whether the loss of *GAL* induction is driven by genomic mutation or by genetic responses. The simultaneous high-level induction of the *INO1*, *NCW2*, and *GRE3* promoters in humulene-producing strains (**Figure 2X-Z**) and the coincidence of amino-acid biosynthetic regulation and *GAL* repression in the FPP accumulating strain (**Supplementary Figure 1 & 2**), are particularly intriguing and lead us to hypothesise that genetic regulation may play a significant role.

With stabilised *GAL* induction, the humulene-producing strains underwent adaptive evolution that improved growth and final titres, albeit at the expense of specific productivity. Proteomic analysis revealed downregulation of the engineered mevalonate-pathway enzymes and upregulation of the humulene biosynthetic modules. These changes indicate that, in addition to humulene tolerance, metabolic balance is critical for stable production; and highlight the importance of designing well-balanced biosynthetic pathways. Overall, this work demonstrates that integrated biocomputing and essential-gene metabolic locking circuits enable the interrogation of non-degenerative evolutionary shifts, providing a valuable platform for metabolic engineering optimisation.

## Materials and Methods

### Strain construction

Plasmids, *Saccharomyces cerevisiae* strains, DNA fragments, and primers used in this study are listed in **Supplementary File 4**. The construction processes for plasmids and yeast strains are also summarised in **Supplementary File 4**.

### Yeast cultivation

Dodecane-overlayed two-phase batch cultivation and semi-fed-batch cultivation were performed to characterise *S. cerevisiae* strains as described previously. Yeast nitrogen base with ammonium sulfate (YNB; FORMEDIUM#CYN0402) buffered with 100 mM 2-(N-morpholino)ethanesulfonic acid (MES) was used as the basal medium [34]. For batch cultivation, MES-buffered 1×YNB (6.9 g L^-1^) and 20 g L^-1^ glucose broth with an initial pH adjusted to 6.0 using ammonium hydroxide were used. For semi-fed-batch cultivation, MES-buffered 4× YNB (27.6 g L^-1^) and 40 g L^-1^ glucose broth with an initial pH adjusted to 5.8 were used [28a]. Yeast cells from glycerol stocks were recovered on 1×YNB-glucose agar supplemented with 125 μM tetracycline. To prepare the seed culture, yeast cells were pre-cultured in YNB-glucose broth without tetracycline for at least 20 h. The seed culture was inoculated to an OD_600_ of approximately 0.2 in 23 ml (for batch cultivation) or 20 ml fresh media (for semi-fed-batch) in 250 ml flasks. In fed-batch cultivation, 400 g L^-1^ glucose stock solution was fed as described above and pH was adjusted by adding 3 μl of 10 M potassium hydroxide per increase in OD_600_. Dodecane (2 ml), sterilised by filtration, was added to sequester terpene products. Yeast cells were sampled subsequently for OD_600_ measurement and flow cytometry analysis. Dodecane was sampled and stored at −20 °C for subsequent qualification.

To measure the promoter activity, cells were pre-cultured in 200 μl 1×YNB (6.9 g L^-1^) and 20 g L^-1^ glucose broth overnight to the stationary phase. Seed culture was then diluted by 1000-fold into fresh broth (1×YNB 20 g L^-1^ glucose) supplemented with estrogen (1 μΜ) or tetracycline (125 μΜ) as described. For the growth in 1×YNB 20 g L^-1^ galactose broth, seed culture was diluted by 100-fold. Cultures were grown to the early exponential phase for flow cytometry analysis.

For plate dropping experiments, yeast cells pre-cultured in YNB-glucose broth with 1 g L^-1^ glutamine supplementation were collected, washed with water, and diluted to OD600 values of 1, 0.1, 0.001, and 0.001. Diluted yeast cells (5 μl) were dropped on the plates and grown for three days before imaging.

### Flow cytometry

Fluorescence of fluorescent reporters in yeast cells was measured by a BD Accuri™ C6 flow cytometry (BD Biosciences, USA) for Figure 1Β or a BD FACSCanto II flow cytometry (Firmware version 1.49) for others. Non-yeast particles were excluded using FSC.H threshold of 250,000 in Accuri C6 flow cytometry and FSC-A (PMT voltage = 200) threshold of 3000 in BD FACSCanto II flow cytometry. For YFAST measurement, 20 μM HMBR, 4-hydroxy-3-methylbenzylidene-rhodanine was added, and for FrFAST measurement, 8 μM tfPoppy, (4-hydroxy-3-methoxy-phenyl)allylidene rhodamine (The Twickle Factory# 555670-250) was added.

In the BD Accuri™ C6 flow cytometry, the yEGFP and YFAST-HMBR were excited by a 488nm laser, and fluorescence was detected through a FL1.A detector equipped with a 530/20 bandpass filter. In the BD FACSCanto II flow cytometry, FITC-A, PerCP-Cy5-5-A, APC-A, and Pacific Blue-A for all detected events were extracted using a BD FACSDiva™ Software (version 8.0.1; CST Version 3.0.1; PLA version 2.0). Fluorescence crosstalk in PerCP-Cy5-5-A channel from green fluorescent protein and Pacific Blue-A channel from E2-Crimson was compensated pragmatically. For each sample, 10,000 events were analysed, and for the samples with a low OD_600_ (<0.5), at least 2000 events were analysed. The fluorescence of each reporter was calculated as fold-changes relative to the background autofluorescence at the corresponding channel in reference to the exponential-phase cells of FP-negative reference strain CEN.PK113-7D.

### Sesquiterpene and monoterpene analysis

Nerolidol, valencene, humulene, farnesol, and limonene were analysed using a HPLC-UV method at the Metabolomics Australia (Queensland Node) as described previously [34]. The dodecane samples that exceeded the analytical linear range were diluted with dodecane. To prepare the samples for HPLC analysis, 5 μl dodecane samples were diluted 40-fold in 195 μl ethanol. The analytes were separated using a Zorbax Extend C18 column (4.6 × 150 mm, 3.5 µm, Agilent PN: 763953-902) equipped with a guard column (SecurityGuard Gemini C18, Phenomenex PN: AJO-7597) at 35 °C. The mobile phase fluid (0.9 ml min-1) contained a mixture of solvent A (water) and solvent B (45% acetonitrile, 45% methanol, and 10% water), with a linear gradient of 5–100% solvent B from 0 to 24 min, then 100% from 24 to 37 min, and finally 5% from 37 to 42 min. Analytes of interest were monitored using a diode array detector (Thermo Fisher Vanquish VF-D11-A) at 202 nm wavelength.

### RNA-sequencing

RNA sequencing was performed using an Oxford Nanopore sequencing platform. Total RNA was extracted from yeast cells using the TRIzol™ Plus RNA Purification Kit (Ambion, catalog #12183555). mRNA was purified using the GenElute™ mRNA Miniprep Kit (Sigma-Aldrich). The PCR-cDNA Barcoding Kit (Oxford Nanopore, catalog #SQK-PCB109) was used to construct the sequencing library. Sequencing of the prepared library was performed on a MinION Mk1C system (MinKNOW version 20.10.6). At least 400 million base pairs of sequencing data were collected per sample. Subsequent sequencing data processing was performed on the Galaxy Australia server [35]. Seqtk seq (Galaxy version 1.5) was used for FASTQ/FASTA format conversion. Minimap2 (Galaxy version) was used to align reads to the reference genome [36]. FeatureCounts (Galaxy version) was used to quantify raw transcript counts for each gene [37]. Transcriptomic data were normalised by scaling the total transcript count per sample to the mean total count across all samples. Gene annotations and functional descriptions were obtained from the *Saccharomyces* Genome Database (SGD: https://www.yeastgenome.org) [38].

### LC-MS/MS Proteomics

Proteomics analysis was performed using a Thermo Fisher Scientific Vanquish-Neo nano UHPLC system coupled with a Thermo Fisher Scientific Orbitrap Exploris 480 Mass Spectrometer and FAIMS Pro-Duo interface at Q-MAP. Proteomics samples were prepared using the S-Trap™ – ProtiFi (Protifi, NY, US) followed manufacture’s protocol. Peptides were separated using a Thermo Fisher Scientific μPAC Neo column C18 (50 cm x 75 μm ID, 100-300 Å) and eluted using a gradient mobile phase mixed with solution A (0.1% formic acid in LCMS grade water) and solution B (0.1% formic acid in LCMS grade 80% aqueous acetonitrile). Full MS scan in positive-ion mode (the electrospray voltage 1.9 kV) was acquired with the m/z range of 340-1110 at resolution of 120,000, automatic gain control (AGC) set to “standard”, and the maximum injection time (MIT) set to “auto”. The MS/MS (DIA) scans were performed with 12 scan events from the mass range of 340-470, 37 scan events from mass range of 460-645, and 10 scan events from mass range of 645-1100, with the mass resolution of 30,000, the AGC target of 5e5 and MIT of 60 ms. Protein abundance data were extracted using Spectronaut v.21.0 (Biognosys AG) via the directDIA⁺ (Deep) workflow against a custom reference proteome database. Search parameters included a Q–value cutoff of ≤ 0.05, at least two quantifying peptides per protein, and factory default settings for false discovery rate (FDR) control, modifications, and retention-time prediction. The transcription factor (TF)–target regulatory matrix was retrieved from the YEASTRACT database (http://www.yeastract.com) [39].

### Statistical analysis

Two-tailed *Welch*’s test was used to calculate *p*-values. Gene-E (version#3.0.215; https://software.broadinstitute.org/GENE-E/) was used for hierarchical clustering analysis using Kendall’s tau distance metric and the average linkage method. Principal Component Analysis was performed using python (version 3.9).

## Data availability

Key plasmids used in this study are available from Addgene (Addgene IDs 223824–223836, 262466–262469, 262471 and 262473). RNA-seq raw reads are available in the Sequence Read Archive (BioProject PRJNA1497633). Proteomics data are available in ProteomeXchange (PXD082605; reviewer access token: dv2uexGgZeUe). The detailed dataset and processed data for the transcriptomic analysis have been deposited in Zenodo (https://zenodo.org/records/22217553). Data, code, and analysis workflows for flow cytometry and proteomics are available on GitHub (https://github.com/pengbingyin/BiocomMetaboLock).

## Acknowledgement

Acknowledgments

B.P. acknowledges insightful discussions with Professor Lars K. Nielsen (The University of Queensland), Dr Robert E. Speight (CSIRO), and Professor Ian Paulsen (Macquarie University). B.P. further acknowledges the mentorship and supervision of Professor Matt Trau during this project and during the CSIRO Future Science Fellowship at The University of Queensland, and the mentorship of Dr Geoff Dumsday at CSIRO. B.P. also acknowledges the supervision and administrative support of Professor Robert E. Speight and Professor Kirill Alexandrov at Queensland University of Technology during this research. B.P. thanks Dr James Antoney (Queensland University of Technology) and Ms Jenny Morcom (Macquarie University) for administrative support associated with the Bioplatforms Australia Omics Research Project Funding Scheme. The authors acknowledge the Queensland Metabolomics and Proteomics (Q-MAP) facility at The University of Queensland for assistance with metabolomics and proteomics data acquisition. Q-MAP is supported by Bioplatforms Australia, an NCRIS-funded initiative. This work was supported by Galaxy Australia, a service provided by Australian BioCommons and its partners. This service receives NCRIS funding through Bioplatforms Australia and The University of Melbourne.

## Funding

This research, and B.P., were supported in part by CSIRO and The University of Queensland through a Synthetic Biology Future Science Platform Fellowship. This research was supported in part by the Australian Government through the Australian Research Council Centres of Excellence funding scheme (project CE200100029). The views expressed herein are those of the authors and are not necessarily those of the Australian Government or the Australian Research Council. This research was also supported in part by Bioplatforms Australia through the Bioplatforms Australia Omics Research Project Funding Scheme, coordinated at Macquarie University.

## Declaration of interests

T.C. is the founder and Chief Executive of Levur, Australia.

## Declaration of generative AI and AI-assisted technologies in the manuscript preparation process

SuperGrok (built by xAI) was used to generate Python codes and check and correct grammar during the preparation of this manuscript. After using this tool, the authors reviewed and edited the content as needed and take full responsibility for the content of the publication.

## Supplemental information

Supplementary file 1: Supplemental information Supplementary file 2: Growth rates

Supplementary file 3: Cell agglomeration of limonene-producing strain in batch cultivation.

Supplementary file 4: Strain, plasmid, and construction processes

## Notes

### Summary of Updates

(1) The abstract was updated to include the narrative on the omics and biosensor methodologies and to improve narrative flow. (2) The second paragraph was revised to provide a clearer overview of the current state of the art in the field. (3) In the Results section, a supplementary video showing the morphological changes in the limonene-producing strain was added. (4) The Discussion was updated to mention previous product-specific biosensor-based metabolic coupling systems, in order to emphasise how the present work differs from earlier studies. (5) Dr John suggested replacing her maiden name with the family name she has used in previous publications. (6) Dr Jahanian updated his affiliation.

## Reference

[1] a) J. Zhang, E. Nuebel, George Q. Daley, Carla M. Koehler, Michael A. Teitell, Cell Stem Cell 2012, 11 (5), 589, 10.1016/j.stem.2012.10.005; b) B. T. Jackson, L. W. S. Finley, Cell Stem Cell 2024, 31 (2), 161, 10.1016/j.stem.2024.01.003; c) J. Zhu, C. B. Thompson, Nature Reviews Molecular Cell Biology 2019, 20 (7), 436, 10.1038/s41580-019-0123-5.

[2] a) C. Danner, R. L. Mach, A. R. Mach-Aigner, Applied Microbiology and Biotechnology 2023, 107 (15), 4745, 10.1007/s00253-023-12615-z; b) G. Wu, Q. Yan, J. A. Jones, Y. J. Tang, S. S. Fong, M. A. G. Koffas, Trends in Biotechnology 2016, 34 (8), 652, 10.1016/j.tibtech.2016.02.010; c) J. Mao, H. Zhang, Y. Chen, L. Wei, J. Liu, J. Nielsen, Y. Chen, N. Xu, Biotechnology Advances 2024, 74, 108401, 10.1016/j.biotechadv.2024.108401; d) I. Menuhin-Gruman, M. Arbel-Groissman, D. Naki, S. Bergman, T. Tuller, Science Advances 11 (40), eadx0796, 10.1126/sciadv.adx0796.

[3] a) B. An, T.-C. Tang, Q. Zhang, T. Wang, Y. Wang, K. Gan, K. Liu, D. L. Zhang, Y. Liu, Y. K. Pan, M. Yu, W. M. Shaw, Q. Liang, Y. Wang, C. A. Vaiana, C. Lou, C. A. Voigt, T. K. Lu, G. M. Church, C. Zhong, Nature 2026, 653 (8114), 587, 10.1038/s41586-026-10259-3; b) H. Doosthosseini, H. Chen, C. A. Voigt, Nature Chemical Biology 2026, 10.1038/s41589-026-02300-3.

[4] a) K. Sechkar, H. Steel, Journal of The Royal Society Interface 2025, 22 (223), 20240602, 10.1098/rsif.2024.0602; b) A. Stone, A. Youssef, S. Rijal, R. Zhang, X.-J. Tian, Trends in Biotechnology 2024, 42 (7), 895, 10.1016/j.tibtech.2024.01.003; c) M. Tartik, ACS Synthetic Biology 2026, 15 (2), 368, 10.1021/acssynbio.5c00715; d) S. M. Bae, M. Chung, J. Y. Lee, S. M. Yoo, Biotechnology Advances 2026, 90, 108889, 10.1016/j.biotechadv.2026.108889.

[5] a) A. Hoynes-O’Connor, T. S. Moon, Current Opinion in Biotechnology 2015, 36, 115, 10.1016/j.copbio.2015.08.007; b) X. Lv, A. Hueso-Gil, X. Bi, Y. Wu, Y. Liu, L. Liu, R. Ledesma-Amaro, Current Opinion in Biotechnology 2022, 76, 102724, 10.1016/j.copbio.2022.102724; c) B. Peng, S. Wei, Journal of Agricultural and Food Chemistry 2025, 73 (17), 10052, 10.1021/acs.jafc.5c01724; d) B. Peng, N. C. Bandari, Z. Lu, C. B. Howard, C. Scott, M. Trau, G. Dumsday, C. E. Vickers, Communications Biology 2022, 5 (1), 135, 10.1038/s42003-022-03070-z; e) B. Peng, M. Tominaga, J. Ishii, E. Marcellin, R. E. Speight, Metabolic Engineering 2026, 102532, 10.1016/j.ymben.2026.102532.

[6] a) C. Fan, H. Chen, Y. Wang, J. Wang, Y. Chen, Y. Zhang, J. Guan, J. Yuan, Advanced Science 2026, 13 (47), e75865, 10.1002/advs.75865; b) L. Y. H. Tan, Z. Lin, J. W. Yeoh, J. Zhang, C. L. Poh, Nature Communications 2026, 17 (1), 6742, 10.1038/s41467-026-73399-0; c) H. M. Beyer, S. Kumar, M. Nieke, C. M. C. Diehl, K. Tang, S. Shumka, C. S. Koh, C. Fleck, J. A. Davies, M. Khammash, M. D. Zurbriggen, Nature Communications 2024, 15 (1), 10470, 10.1038/s41467-024-54350-7.

[7] H. Zhai, X. Liu, P. Guo, Y. Guo, X. Yang, Q. Qi, J. Nielsen, Z. Liu, Y. Chen, J. Hou, Trends in Biotechnology, 10.1016/j.tibtech.2026.04.007.

[8] F. Meng, W. M. Shaw, Y. K. K. Kam, T. Ellis, Cell 2025, 188 (18), 4936, 10.1016/j.cell.2025.06.025.

[9] a) B. D. Huang, Y. Yu, J. Lee, M. W. Repasky, Y. Xie, M. J. Realff, C. J. Wilson, Nature Communications 2025, 16 (1), 9414, 10.1038/s41467-025-64457-0; b) W. Cao, L. Liu, Q. Sun, Y. Shan, Y. Chen, Nature Communications 2025, 16 (1), 7037, 10.1038/s41467-025-62464-9; c) N. Kim, G. De Carluccio, K. Zhang, J. J. Collins, Cell Systems 2026, 17 (2), 101533, 10.1016/j.cels.2026.101533.

[10] a) M. D. J. Bragdon, N. Patel, J. Chuang, E. Levien, C. J. Bashor, A. S. Khalil, Cell 2023, 186 (18), 3810, 10.1016/j.cell.2023.07.012; b) J. L. Chlebek, S. P. Leonard, C. Kang-Yun, M. C. Yung, D. P. Ricci, Y. Jiao, D. M. Park, Nucleic Acids Research 2023, 51 (13), 7094, 10.1093/nar/gkad484; c) D. P. Byrom, A. P. S. Darlington, Nature Communications 2025, 16 (1), 8590, 10.1038/s41467-025-63627-4.

[11] R. H. Dahl, F. Zhang, J. Alonso-Gutierrez, E. Baidoo, T. S. Batth, A. M. Redding-Johanson, C. J. Petzold, A. Mukhopadhyay, T. S. Lee, P. D. Adams, J. D. Keasling, Nature Biotechnology 2013, 31 (11), 1039, 10.1038/nbt.2689.

[12] a) J. Savigny, K. Shabestary, M. Portela, C. Klemm, Y. Sum, P. Hapeta, M. Storch, C. Rowlands, R. Ledesma-Amaro, Nature Communications 2025, 17 (1), 645, 10.1038/s41467-025-67408-x; b) L. Qin, B. Yang, R. Huang, H. Zhang, Q. Li, X. Li, Y. Tan, W. Sun, H. Jia, B. Hu, B. Lv, C. Li, Nature Communications 2026, 17 (1), 5081, 10.1038/s41467-026-71801-5.

[13] Z. Ye, Y. Huang, B. Shi, Z. Xiang, Z. Tian, M. Huang, L. Wu, Z. Deng, K. Shen, T. Liu, Metabolic Engineering 2022, 72, 107, 10.1016/j.ymben.2022.03.005.

[14] a) B. Peng, A. Jahanian, T. R. Collier, H. Peng, J. B. Behrendorff, R. E. Speight, Food Chemistry 2026, 503, 147749, 10.1016/j.foodchem.2025.147749; b) Y. Yu, X. Fu, J. Hu, J. Nielsen, S. Shi, Current Opinion in Biotechnology 2025, 96, 103387, 10.1016/j.copbio.2025.103387.

[15] a) D. Canadell, N. Ortiz-Vaquerizas, S. Mogas-Diez, E. de Nadal, J. Macia, F. Posas, Nucleic Acids Research 2022, 50 (21), 12578, 10.1093/nar/gkac1120; b) Y. Chen, S. Zhang, E. M. Young, T. S. Jones, D. Densmore, C. A. Voigt, Nature Microbiology 2020, 5 (11), 1349, 10.1038/s41564-020-0757-2.

[16] a) R. Balakrishnan, J. Park, K. Karra, B. C. Hitz, G. Binkley, E. L. Hong, J. Sullivan, G. Micklem, J. Michael Cherry, Database 2012, 2012, bar062, 10.1093/database/bar062; b) W. Chen, M. Han, J. Zhou, Q. Ge, P. Wang, X. Zhang, S. Zhu, L. Song, Y. Yuan, National Science Review 2021, 8 (5), nwab028, 10.1093/nsr/nwab028.

[17] a) R. S. K. Walker, E. Archer, P. Erpf, I. T. Paulsen, I. S. Pretorius, FEMS Yeast Research 2026, 26, foag034, 10.1093/femsyr/foag034; b) H. Wang, J. Nielsen, Y. J. Zhou, H. Lu, Proceedings of the National Academy of Sciences 2025, 122 (32), e2502044122, doi:10.1073/pnas.2502044122.

[18] a) B. Peng, L. K. Nielsen, S. C. Kampranis, C. E. Vickers, Metabolic Engineering 2018, 47, 83, 10.1016/j.ymben.2018.02.005; b) L. Chen, W. Xiao, M. Yao, Y. Wang, Y. Yuan, Frontiers in Bioengineering and Biotechnology 2023, Volume 11 - 2023, 10.3389/fbioe.2023.1132244.

[19] I. F. Hayat, M. Plan, B. E. Ebert, G. Dumsday, C. E. Vickers, B. Peng, Microbial Biotechnology 2021, 14 (6), 2627, 10.1111/1751-7915.13880.

[20] M.-A. Plamont, E. Billon-Denis, S. Maurin, C. Gauron, F. M. Pimenta, C. G. Specht, J. Shi, J. Quérard, B. Pan, J. Rossignol, K. Moncoq, N. Morellet, M. Volovitch, E. Lescop, Y. Chen, A. Triller, S. Vriz, T. Le Saux, L. Jullien, A. Gautier, Proceedings of the National Academy of Sciences 2016, 113 (3), 497, 10.1073/pnas.1513094113.

[21] Z. Lu, S. Evans, L. McDonnell, N. C. Bandari, Y. Weng, W. Jin, R. Speight, G. Schenk, C. B. Howard, C. E. Vickers, B. Peng, Yeast 2025, 42 (8-10), 169, 10.1002/yea.4005.

[22] D. E. Martin, A. Soulard, M. N. Hall, Cell 2004, 119 (7), 969, 10.1016/j.cell.2004.11.047.

[23] A. Komeili, K. P. Wedaman, E. K. O’Shea, T. Powers, Journal of Cell Biology 2000, 151 (4), 863, 10.1083/jcb.151.4.863.

[24] S. M. O’Rourke, I. Herskowitz, E. K. O’Shea, Trends in Genetics 2002, 18 (8), 405, 10.1016/S0168-9525(02)02723-3.

[25] S. Evans, M. Tominaga, Z. Lu, N. C. Bandari, L. McDonnell, C. Wang, R. E. Speight, J. Ishii, C. E. Vickers, B. Peng, ACS Synthetic Biology 2025, 14 (9), 3804, 10.1021/acssynbio.5c00432.

[26] B. Peng, M. R. Plan, P. Chrysanthopoulos, M. P. Hodson, L. K. Nielsen, C. E. Vickers, Metabolic Engineering 2017, 39, 209, 10.1016/j.ymben.2016.12.003.

[27] J. Yang, M. Xian, S. Su, G. Zhao, Q. Nie, X. Jiang, Y. Zheng, W. Liu, *PLOS ONE* 2012, *7* (4), e33509, 10.1371/journal.pone.0033509.

[28] a) Z. Lu, Q. Shen, N. C. Bandari, S. Evans, L. McDonnell, L. Liu, W. Jin, Carlos H. Luna-Flores, T. Collier, G. Talbo, T. McCubbin, L. Esquirol, C. Myers, M. Trau, G. Dumsday, R. Speight, Christopher B. Howard, Claudia E. Vickers, B. Peng, Nucleic Acids Research 2024, 52 (12), 7367, 10.1093/nar/gkae460; b) B. Peng, L. Esquirol, Z. Lu, Q. Shen, L. C. Cheah, C. B. Howard, C. Scott, M. Trau, G. Dumsday, C. E. Vickers, Nature Communications 2022, 13 (1), 2895, 10.1038/s41467-022-30529-8; c) B. Peng, S. J. Weintraub, Z. Lu, S. Evans, Q. Shen, L. McDonnell, M. Plan, T. Collier, L. C. Cheah, L. Ji, C. B. Howard, W. Anderson, M. Trau, G. Dumsday, E. L. Bredeweg, E. M. Young, R. Speight, C. E. Vickers, ACS Synthetic Biology 2023, 13 (1), 141, 10.1021/acssynbio.3c00363.

[29] Z. Lu, B. Peng, B. E. Ebert, G. Dumsday, C. E. Vickers, Nature Communications 2021, 12 (1), 1051, 10.1038/s41467-021-21313-1.

[30] H. C. Yoo, Y. C. Yu, Y. Sung, J. M. Han, Experimental & Molecular Medicine 2020, 52 (9), 1496, 10.1038/s12276-020-00504-8.

[31] L. McDonnell, S. Evans, Z. Lu, M. Suchoronczak, J. Leighton, E. Ordeniza, B. Ritchie, N. Valado, N. Walsh, J. Antoney, C. Wang, C. H. Luna-Flores, C. Scott, R. Speight, C. E. Vickers, B. Peng, Synthetic and Systems Biotechnology 2024, 9 (4), 820, 10.1016/j.synbio.2024.06.009.

[32] a) T. Jordá, S. Puig, Regulation of Ergosterol Biosynthesis in Saccharomyces cerevisiae. In Genes, 2020; Vol. 11, p 795; b) M. A. Wangeline, R. Y. Hampton, J Biol Chem 2018, 293 (38), 14937, 10.1074/jbc.RA118.001808.

[33] a) V. D’Ambrosio, E. Dore, R. Di Blasi, M. van den Broek, S. Sudarsan, J. t. Horst, F. Ambri, M. O. A. Sommer, P. Rugbjerg, J. D. Keasling, R. Mans, M. K. Jensen, Metabolic Engineering 2020, 61, 369, 10.1016/j.ymben.2020.07.006; b) Y. Lv, Y. Gu, J. Xu, J. Zhou, P. Xu, Metabolic Engineering 2020, 61, 79, 10.1016/j.ymben.2020.05.005.

[34] B. Peng, M. R. Plan, A. Carpenter, L. K. Nielsen, C. E. Vickers, Biotechnology for Biofuels 2017, 10 (1), 43, 10.1186/s13068-017-0728-x.

[35] C. The Galaxy, Nucleic Acids Research 2026, 54 (W1), W105, 10.1093/nar/gkag469.

[36] H. Li, Bioinformatics 2018, 34 (18), 3094, 10.1093/bioinformatics/bty191.

[37] Y. Liao, G. K. Smyth, W. Shi, Bioinformatics 2014, 30 (7), 923, 10.1093/bioinformatics/btt656.

[38] S. R. Engel, S. Aleksander, R. S. Nash, E. D. Wong, S. Weng, S. R. Miyasato, G. Sherlock, J. M. Cherry, Genetics 2025, 229 (3), iyae185, 10.1093/genetics/iyae185.

[39] M. C. Teixeira, R. Viana, M. Palma, J. Oliveira, M. Galocha, M. N. Mota, D. Couceiro, M. G. Pereira, M. Antunes, I. V. Costa, P. Pais, C. Parada, C. Chaouiya, I. Sá-Correia, P. T. Monteiro, Nucleic Acids Research 2023, 51 (D1), D785, 10.1093/nar/gkac1041.

