## Supplementary file 1 for "Evolutionary stabilisation of stressful metabolism *via* integrated biocomputing and essential-gene metabolic locking circuits"

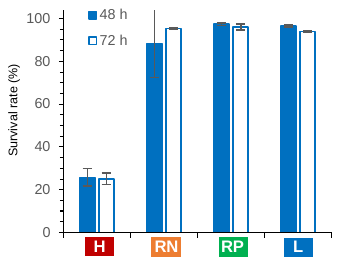

Supplementary Figure 1. Survival rates of the four strains in batch cultivation in Figure 1B: H (ILHA oG57BRB), RN (ILHA oG57H2BRB), RP (ILHA NG57BRB), and L (ILHA NG57H2BRB). Mean values are shown (n = 3 biological replicates).

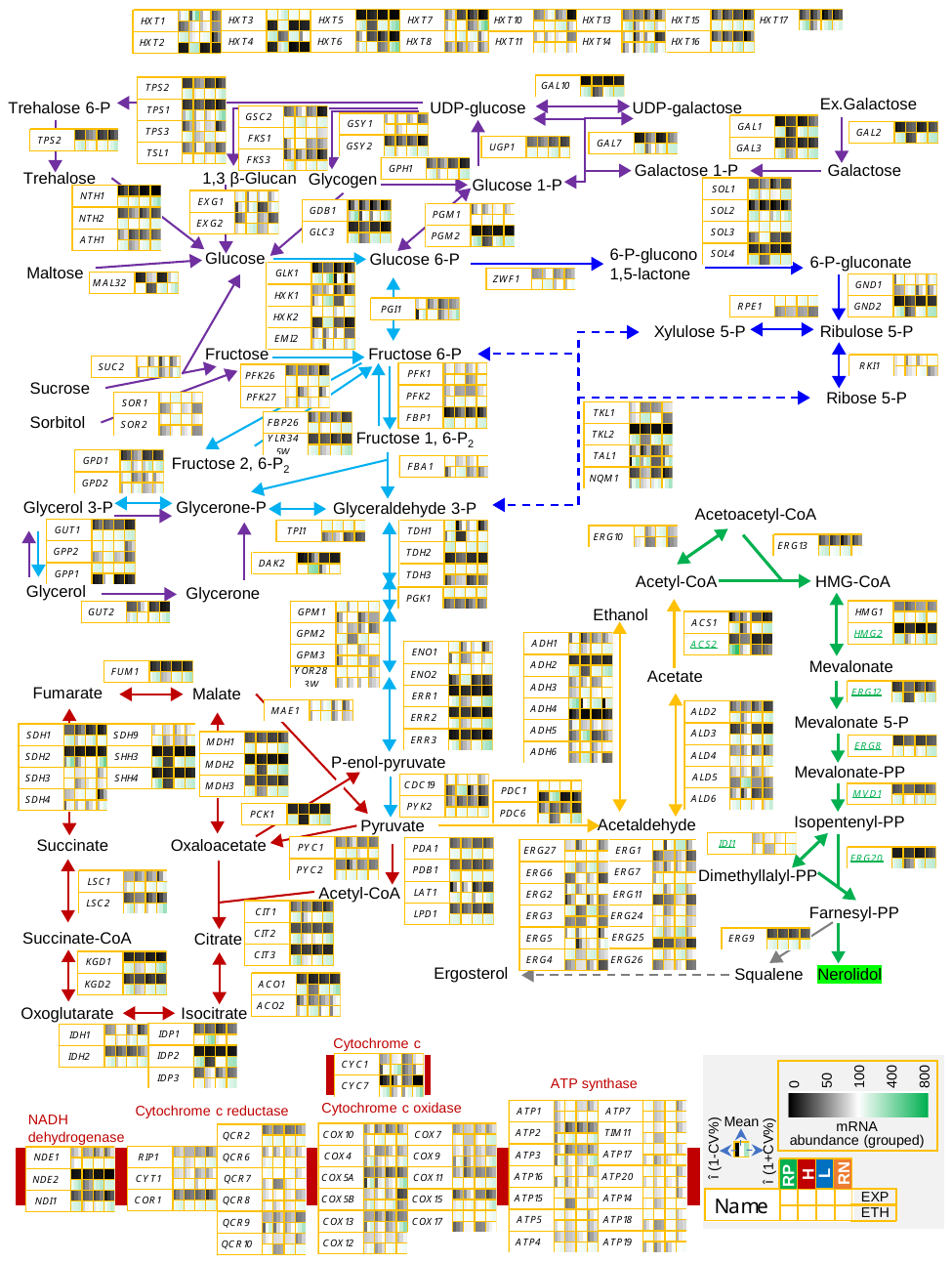

Supplementary Figure 2. Transcriptomic analysis of carbon hydrate catabolic pathways in strains RP (ILHA NG57BRB), H (ILHA oNG57BRB), L (ILHA NG57BH2BRB), and RN (ILHA oG57BH2BRB). Referring to Figure 1.

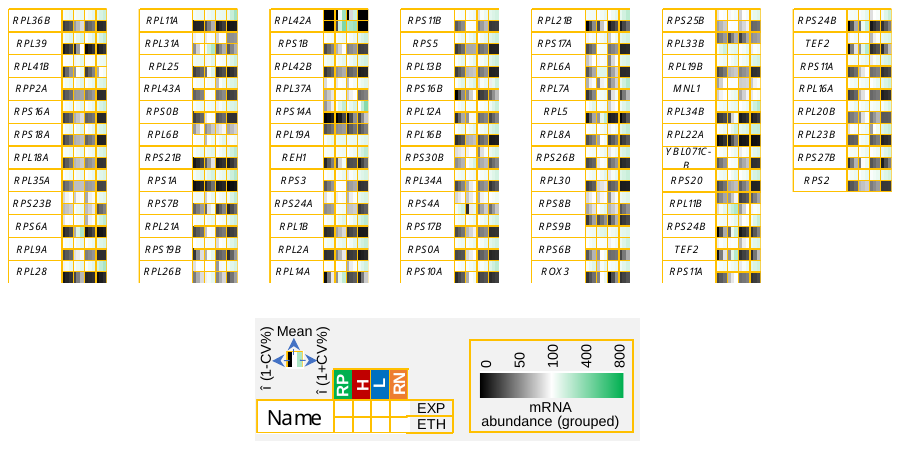

Supplementary Figure 3. Transcriptomic analysis of ribosome components in strains RP (ILHA NG57BRB), H (ILHA oNG57BRB), L (ILHA NG57BH2BRB), and RN (ILHA oG57BH2BRB). Referring to Figure 1.

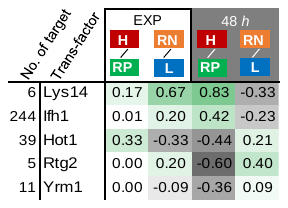

Supplementary Figure 4. Transcription factor enrichment analysis for RNA-seq comparisons among strains RP (ILHA NG57BRB), H (ILHA oNG57BRB), L (ILHA NG57BH2BRB), and RN (ILHA oG57BH2BRB). Genes with an absolute fold-change ≥ 0.7 and a two-tailed *Welch*’s *t*-test *p*-value ≤ 0.1 were classified as up- or downregulated in each comparison. Enrichment values were calculated as the difference (Δ) between the ratios of upregulated and downregulated target genes controlled by each transcription factor. Only transcription factors with an absolute enrichment value ≥ 0.3 and at least five regulated target genes are shown. Referring to Figure 1.

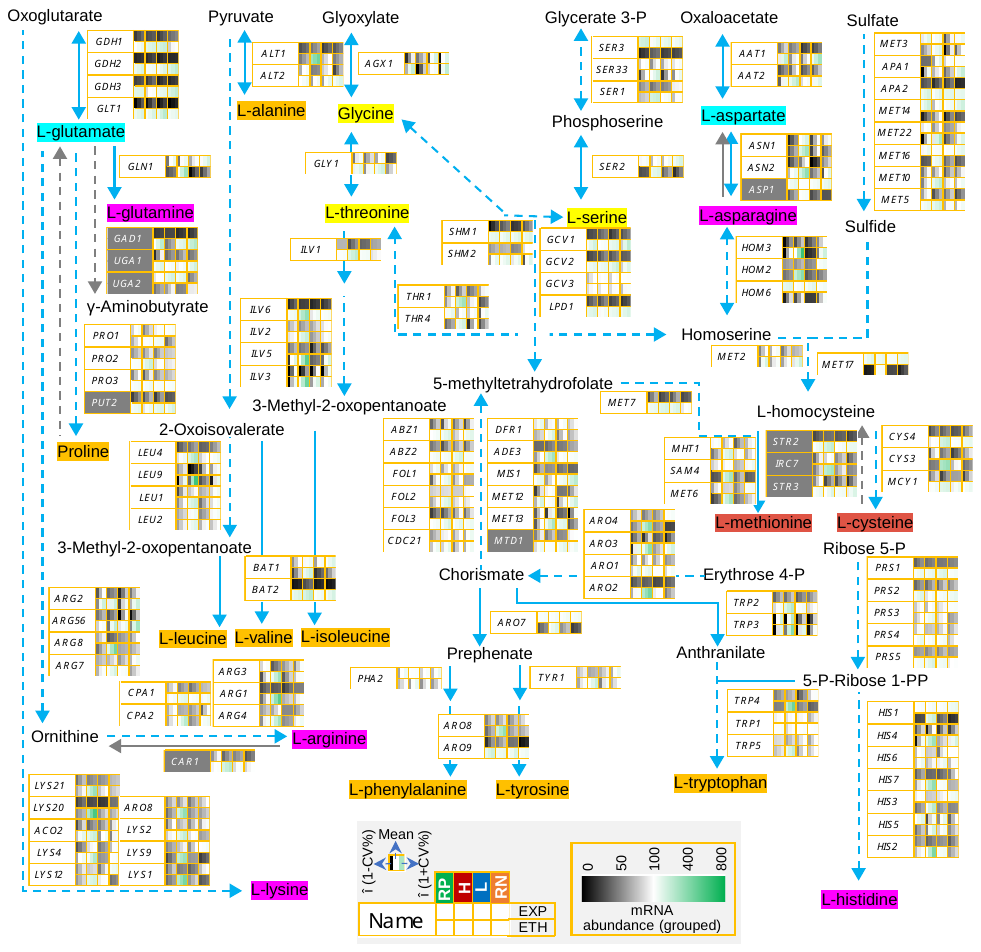

Supplementary Figure 5. Transcriptomic analysis of amino acid biosynthetic pathways in strains RP (NG57BRB), H (oNG57BRB), L (NG57BH2BRB), and RN (oG57BH2BRB). Referring to Figure 1.

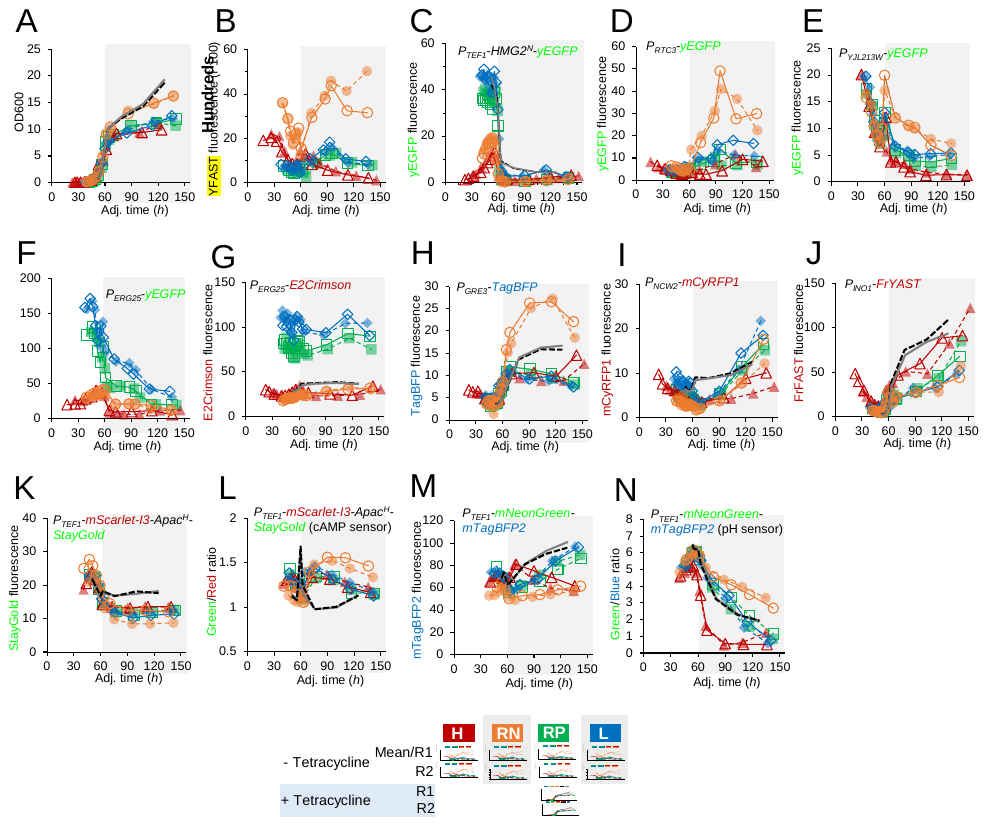

Supplementary Figure 6. Characterisation of the activity of promoters and the response of cAMP- and pH-biosensors in strains RP (NG57BRB), H (oNG57BRB), L (NG57BH2BRB), and RN (oG57BH2BRB). Seed cells were inoculated into medium supplemented with 1 mM 1-naphthaleneacetic acid to an OD_600_ of 0.001 and grown overnight before measurements were started. Time was adjusted by aligning the point at which OD_600_ = 7 with 60 h on the x-axis. YNB 20 g L^-1^ glucose medium buffered with 100 mM MES-NH4^+^ (pH = 6) was used. Tetracycline was added at the concentration of 125 mM. The mean values (N ≥ 3) were shown in A, and two biological replicates (R1 & R2) was characterised in B-N.

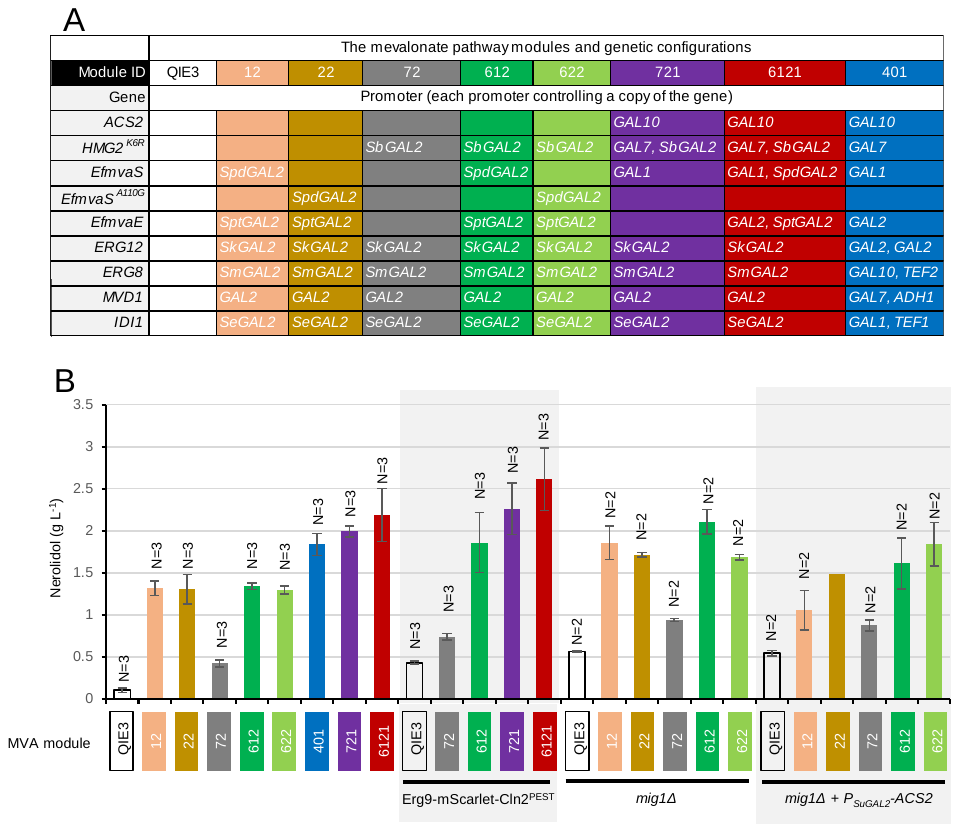

Supplementary Figure 7. Characterisation of metabolic pathway engineering modules for nerolidol production. (A) Configuration of the mevalonate pathway engineering modules. (B) Nerolidol production at 72 h in two-phase batch cultivation in MES-buffered YNB broth with 20 g L^-1^ glucose as the carbon source. Mean values ± standard deviation (N = 3) or Mean values ± absolute error (N = 2) are shown.

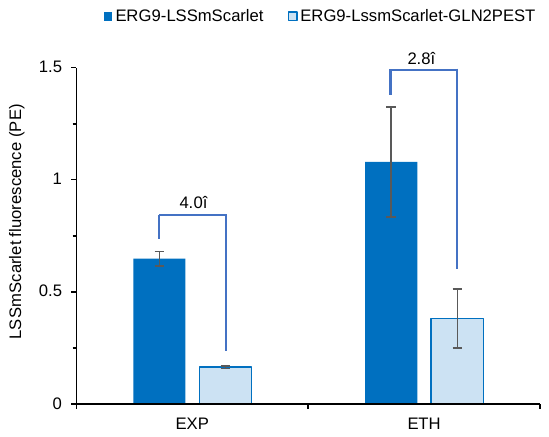

Supplementary Figure 8. Comparing LSSmScarlet fluorescence in the strains with or without Cln2 PEST sequence fused to the C-terminus of squalene synthase Erg9 in CEN.PK113-7D. EXP, the exponential growth phase (OD_600_ < 2); ETH, the ethanol growth phase (OD_600_ > 10). YNB 20 g L^-1^ glucose medium buffered with 100 mM MES-NH4^+^ (pH = 6) was used. Mean values ± standard deviation are shown (n = 3 biological replicates).

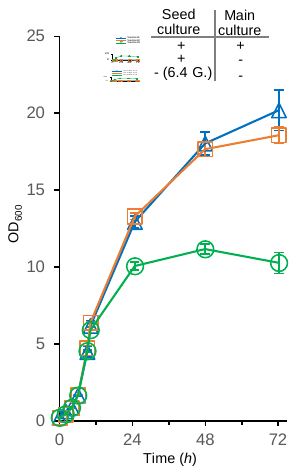

Supplementary Figure 9. Growth profiles for batch cultivation in Figure 2C.

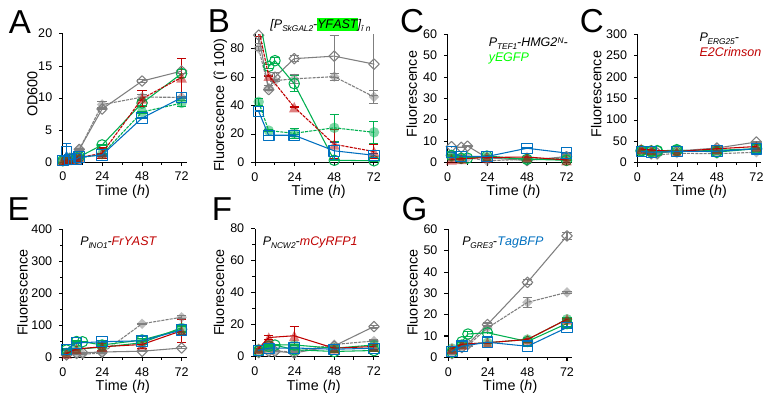

Supplementary Figure 10. Characterisation of the strains equipped with six versions of prenyl pyrophosphate biosynthetic modules and lacking heterologous terpene-synthetic modules.

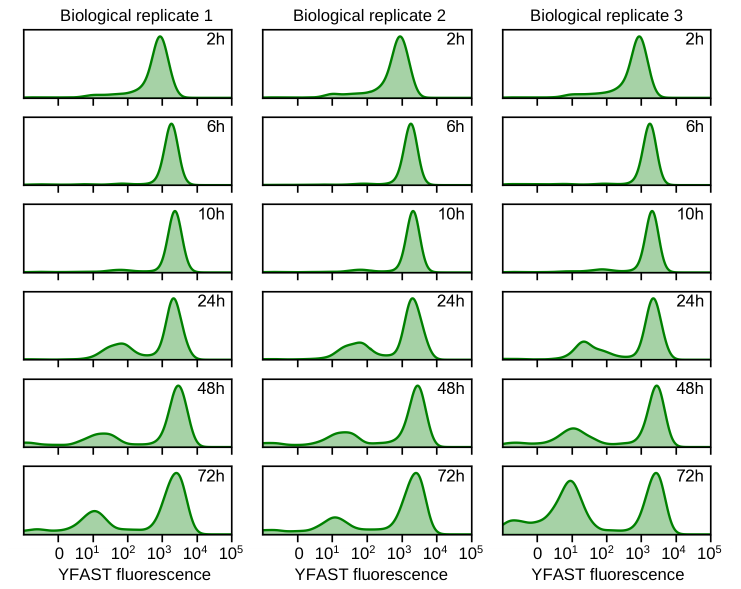

Supplementary Figure 11. Density distribution of YFAST fluorescence in nerolidol-producing #612D strain (ILHA NPME612D). Referring to Figure 2E.

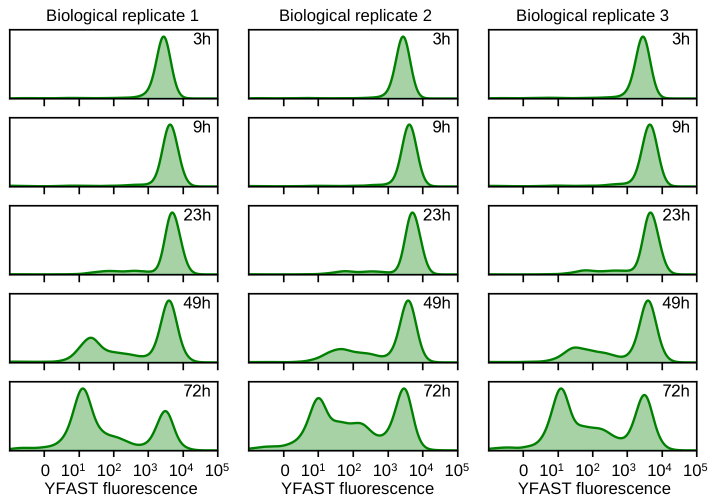

Supplementary Figure 12. Density distribution of YFAST fluorescence in valencene-producing #612D strain (ILHA VLPME612D). Referring to Figure 2M.

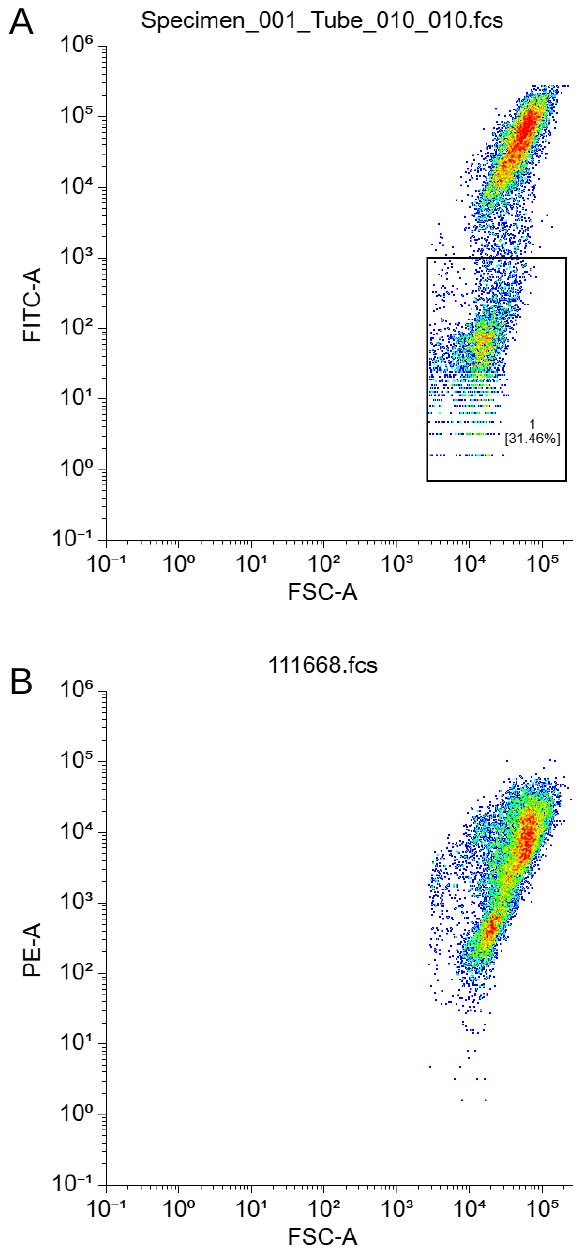

Supplementary Figure 13. Calculation of the ratio of cells losing YFAST fluorescence (A) and propidium iodide (B) in the nerolidol-producing #612D strain (ILHA NPMED612; biological replicate 1) at 72 *h*. Data were visualised using <https://floreada.io/analysis>.

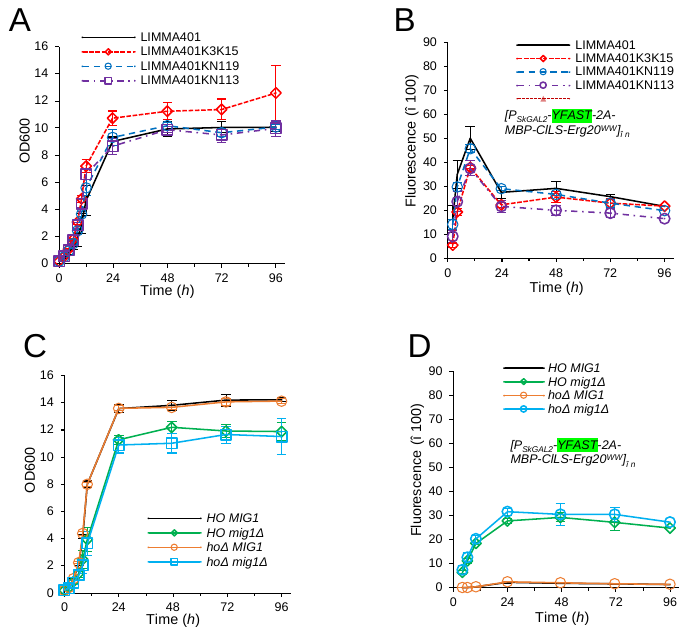

Supplementary Figure 14. Growth and YFAST fluorescence profiles for batch cultivation of the strains in Figure 3B.

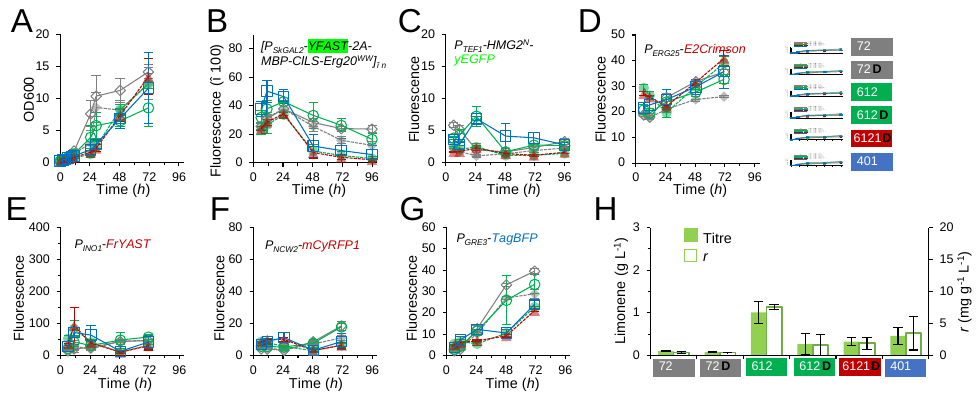

Supplementary Figure 15. Characterisation of limonene-producing strains in *ERG20*-wildtype background. Supplementary to Figure 3.

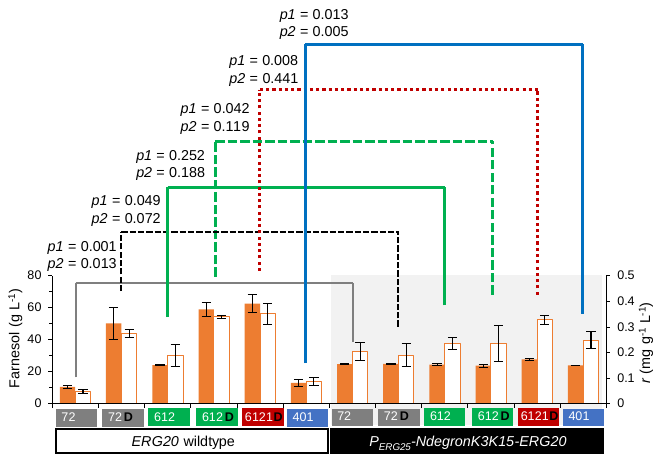

Supplementary Figure 16. Farnesol production in limonene-producing strains at 96 *h*. Supplementary data for Figure 3C-J and Supplementary Figure 15. Two-tailed *Welch*’s test was used to calculate *p* value: *p1*, titre; and *p2*, rate.

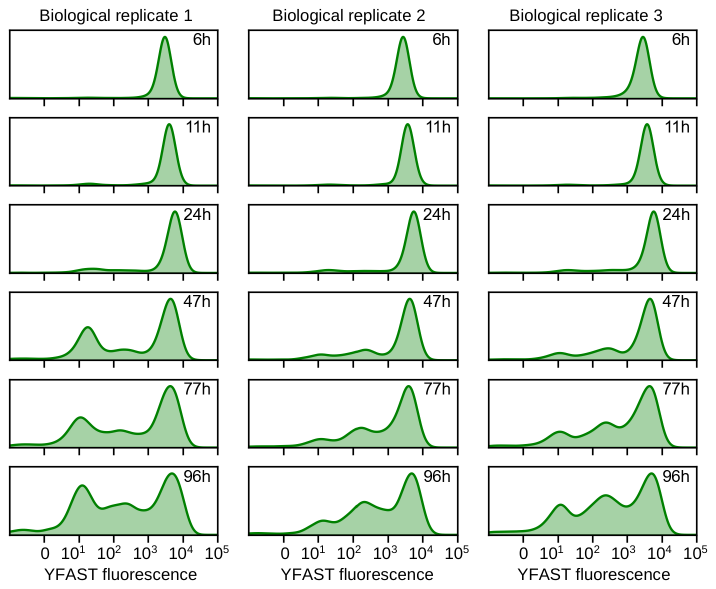

Supplementary Figure 17. Density distribution of YFAST fluorescence in valencene-producing #612 strain (ILHA LIMPME612K3K15). Referring to Figure 3D.

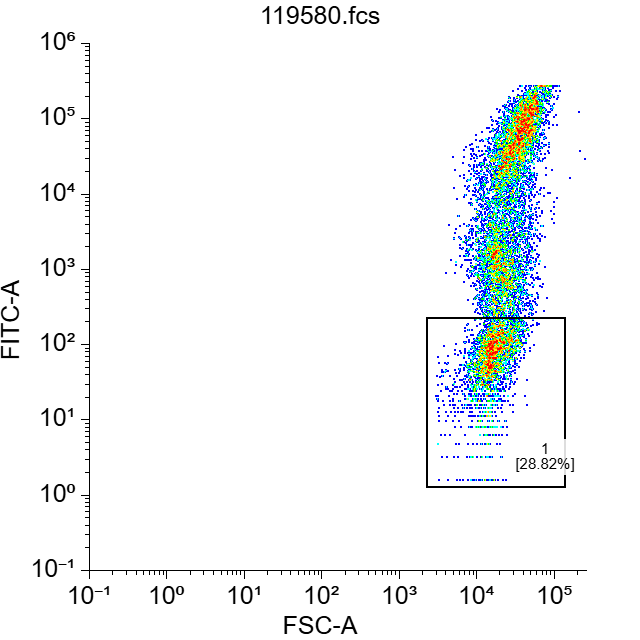

Supplementary Figure 18. Example of flow cytometry data to evaluate the population that lost YFAST fluorescence in limonene-producing #612 strain (ILHA LIMPME612K3K15) at 96 *h*. Data were visualised using <https://floreada.io/analysis>.

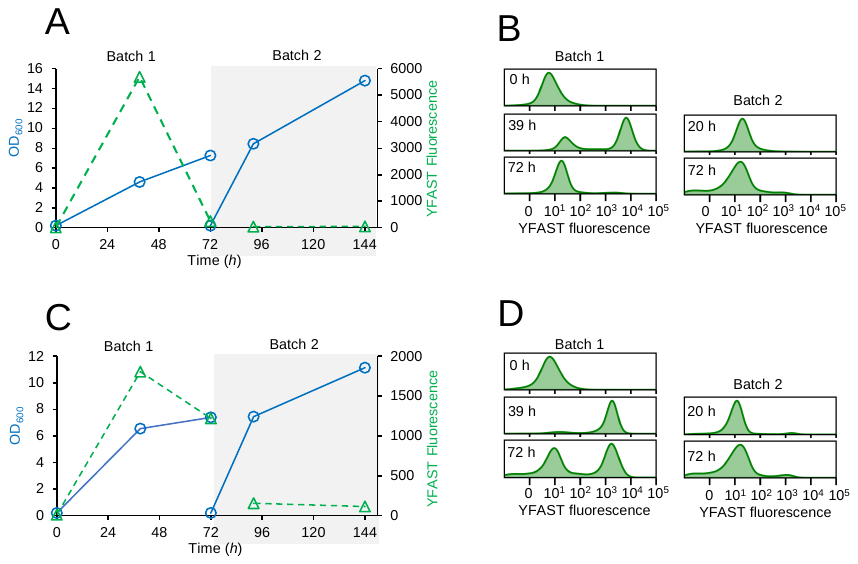

Supplementary Figure 19. Transferring cultivation of NPME612D, and VLPME612D.

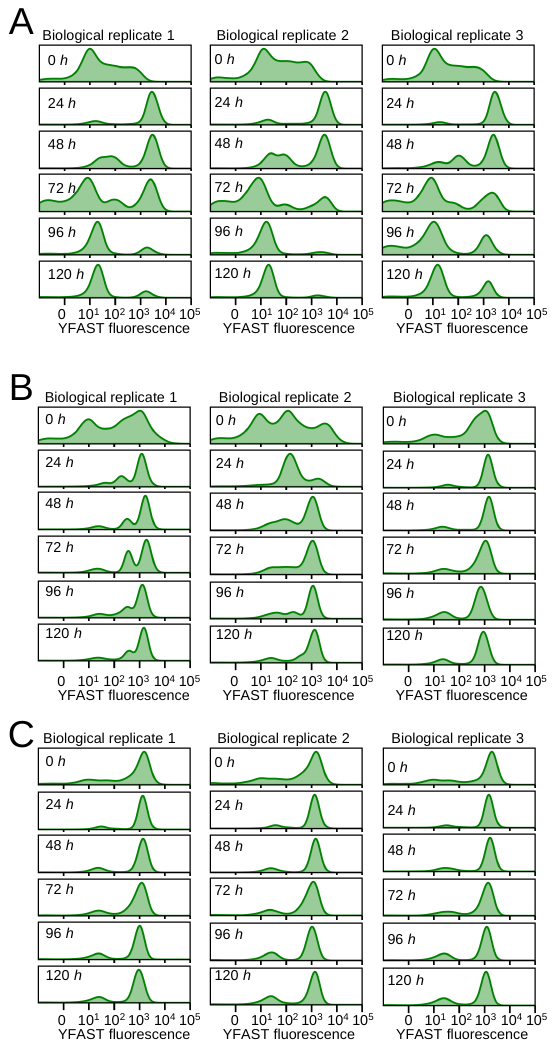

Supplementary Figure 20. Density distribution of YFAST fluorescence in nerolidol-producing strain (NPMER612Dg) in fed-batch cultivation. Referring to Figure 5D: ILHA NPME612D (A), ILHA NPMER612Dg (B), and ILHA NPMER612DgE (C).

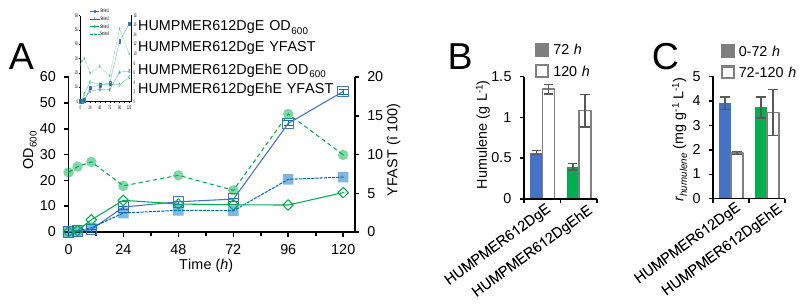

Supplementary Figure 21. Characterisation of adapted humulene-producing strains in the cultivation with glucose feeding at 72 *h*. The MES-buffered 4×YNB broth with 40 g L^-1^ glucose as the initial carbon source, with 40 g L^-1^ glucose and 1×YP (20 g L^-1^ peptone, 10 g L^-1^ yeast extract) fed at 72 *h*.

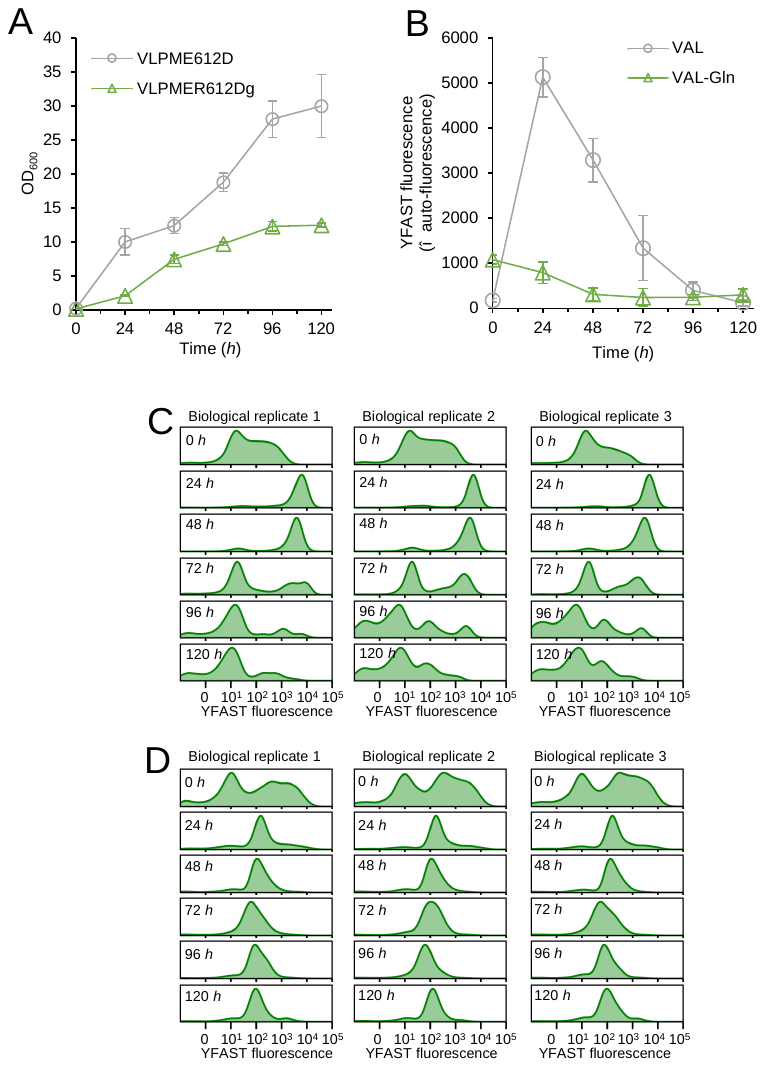

Supplementary Figure 22. Growth and YFAST fluorescence profiles of valencene-producing strains ILHA VLPME612D and ILHA VLPMER612D (the redundant and complementation module and the metabolic locking module) in fed-batch cultivation. (A) Growth. (B) YFAST fluorescence. (C) Density distribution of YFAST fluorescence in ILHA VLPME612D. (D) Density distribution of YFAST fluorescence in ILHA VLPMER612Dg. The MES-buffered 4×YNB broth with 40 g L^-1^ glucose as the initial carbon source, with 40 g L^-1^ glucose fed at 72 and 96 *h*.

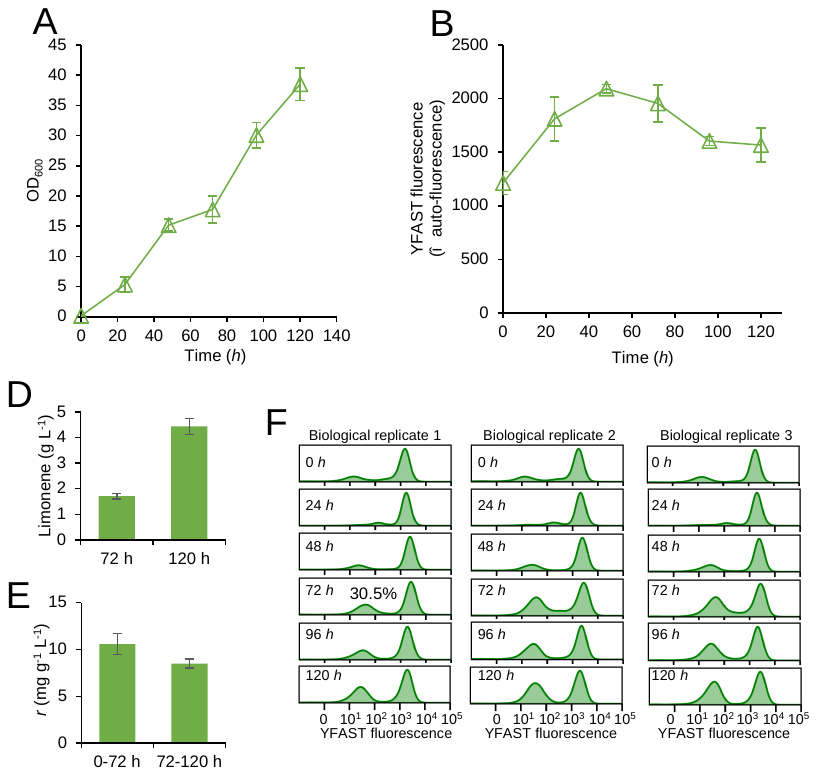

Supplementary Figure 23. Semi-fed-batch characterisation of the limonene-producing strain carrying the *GLN1*-mediated locking mechanism. (A) Growth. (B) YFAST fluorescence. (D) Limonene titres. (E) Specific limonene production rates. (F) Density distribution of YFAST fluorescence. The MES-buffered 4×YNB broth with 40 g L^-1^ glucose as the initial carbon source. Glucose (40 g L^-1^) was fed at 72 *h* and 96 *h*.

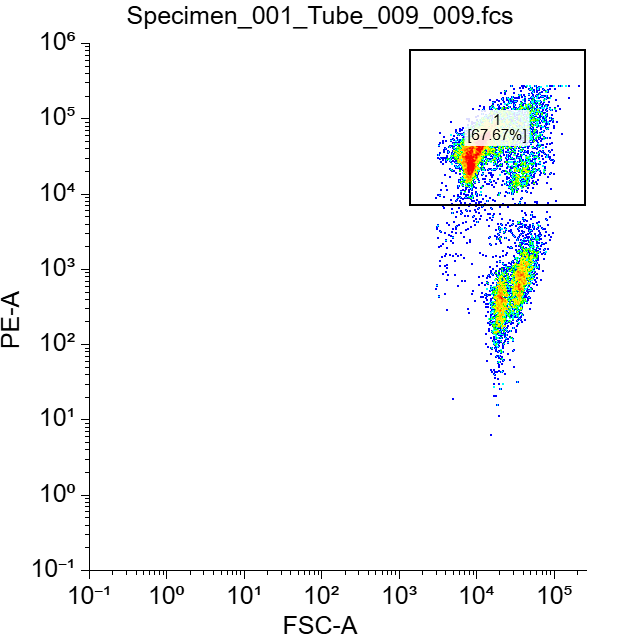

Supplementary Figure 24. Example of propidium iodide in adapted limonene-producing strain LIMPMER612K3K15g.

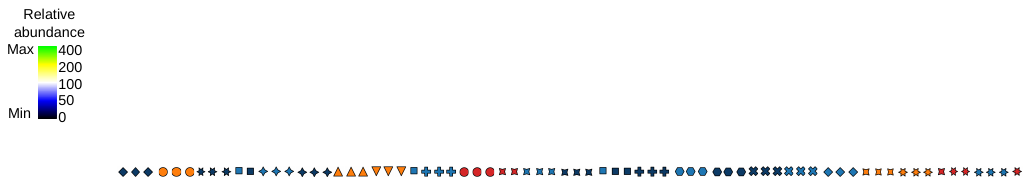

Supplementary Figure 25. Clustering analysis of proteomics data in nerolidol-, valencene-, humulene-, and limonene-producing strains: CENPK (CEN.PK113-7D), NER (ILHA NPME612D), NERg (ILHA NPMER612Dg), NERgE (ILHA NPMER612DE), VAL (ILHA VLPME612D), VALg (ILHA VLPMER612Dg), HUMg (ILHA HUMPMER612Dg), HUMgE (ILHA HUMPMER612DgE), HUMgEh (ILHA HUMPMER612DgEh), HUMgEhE (ILHA HUMPMER612DgEhE), and LIMg (ILHA LIMPMER612K3K15g).

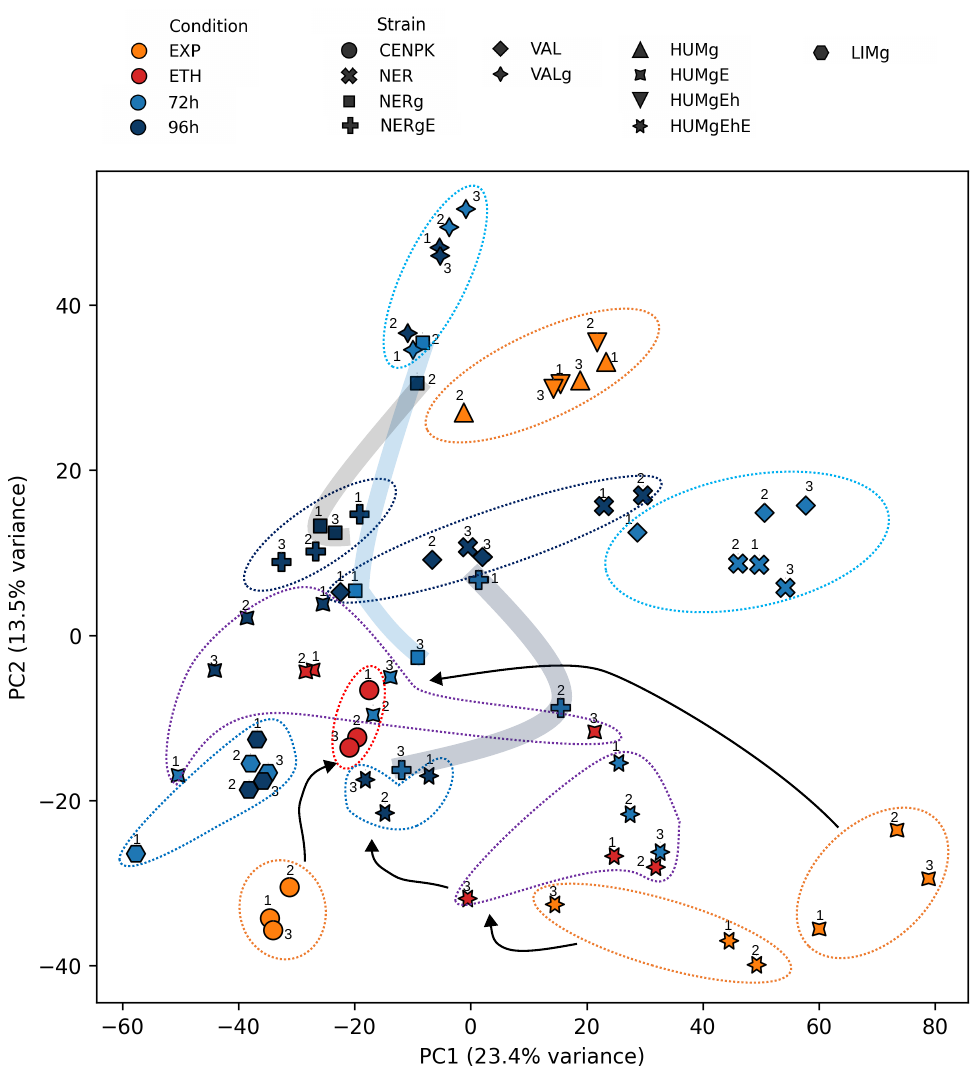

Supplementary Figure 26. Principal Component Analysis (PCA) analysis of proteomics data in nerolidol-, valencene-, humulene-, and limonene-producing strains: CENPK (CEN.PK113-7D), NER (ILHA NPME612D), NERg (ILHA NPMER612Dg), NERgE (ILHA NPMER612DE), VAL (ILHA VLPME612D), VALg (ILHA VLPMER612Dg), HUMg (ILHA HUMPMER612Dg), HUMgE (ILHA HUMPMER612DgE), HUMgEh (ILHA HUMPMER612DgEh), HUMgEhE (ILHA HUMPMER612DgEhE), and LIMg (ILHA LIMPMER612K3K15g).

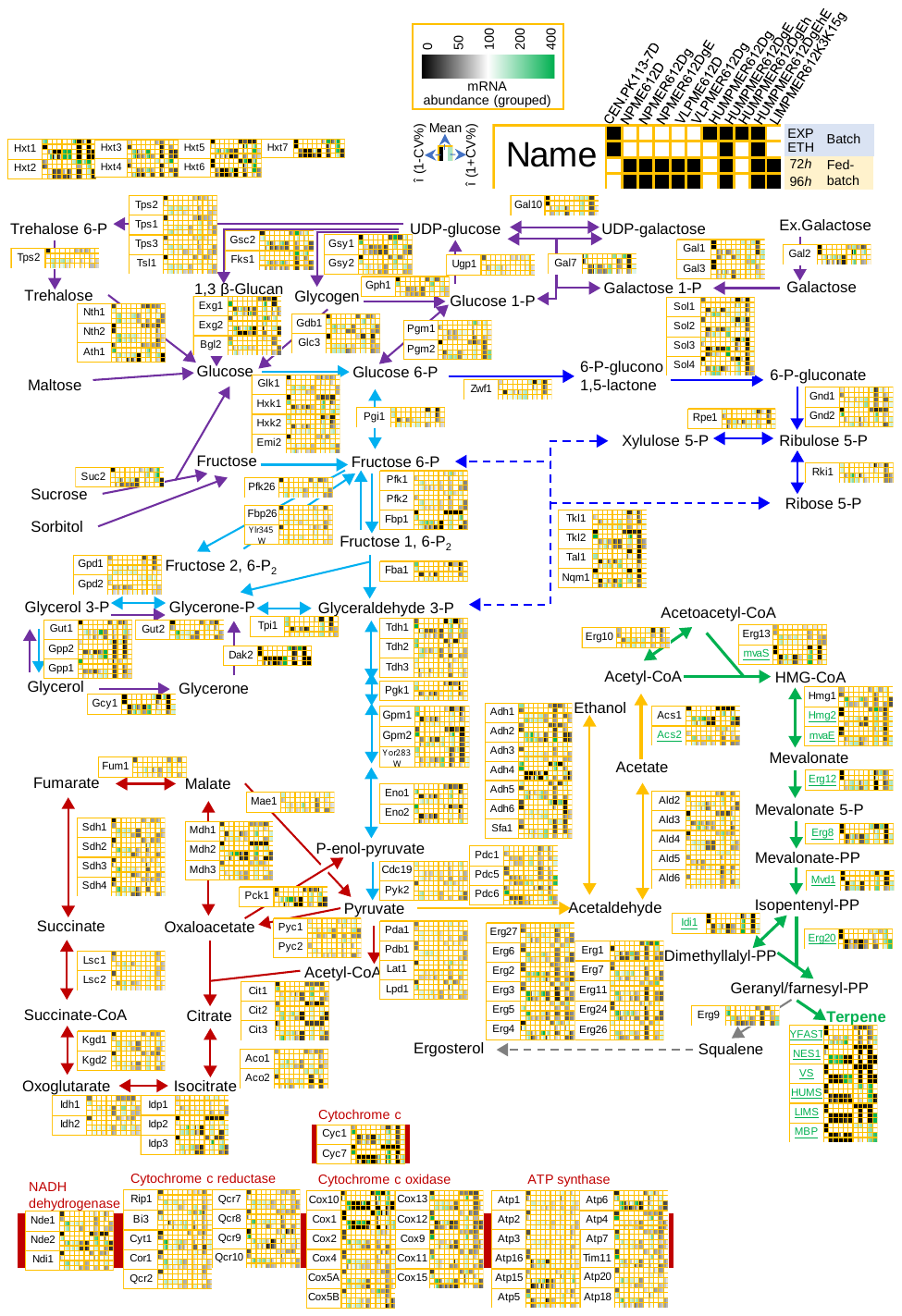

Supplementary Figure 27. Proteomics analysis of carbon hydrate catabolic pathways in nerolidol-, valencene-, humulene-, and limonene-producing strains.

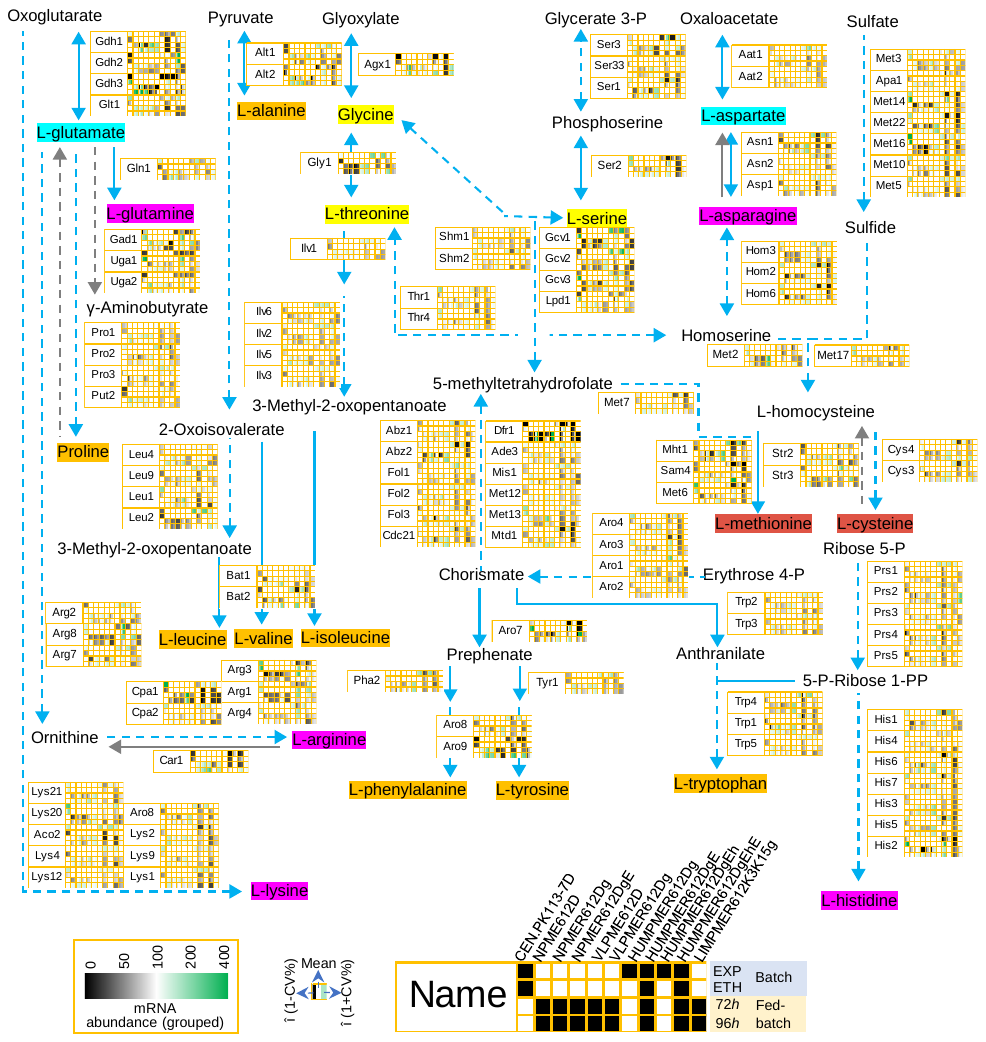

Supplementary Figure 28. Proteomics analysis of amino acid biosynthetic pathways in nerolidol-, valencene-, humulene-, and limonene-producing strains.
